# Cognitive and neural bases of decision-making causing civilian casualties during intergroup conflict

**DOI:** 10.1101/2020.12.04.411280

**Authors:** Xiaochun Han, Shuai Zhou, Nardine Fahoum, Taoyu Wu, Tianyu Gao, Simone Shamay-Tsoory, Michele J. Gelfand, Xinhuai Wu, Shihui Han

## Abstract

Civilian casualties occur during military attacks. Such “collateral damage” is prohibited by international laws but increases with substantial consequences when intergroup conflict escalates. Here, we investigate cognitive and neural bases of decision-making processes resulting in civilian harm, using a task that simulates punishment decision-making during intergroup conflict. We test two groups of Chinese participants in a laboratory setting, and two ethnic groups (Jewish and Palestinian) in Israel. The results dissociate two psychological constructs, harm preference and harm avoidance, which respectively characterize punishment decision-making related to outgroup combatants and outgroup noncombatants during intergroup conflict. In particular, individuals show decreased avoidance of harming outgroup noncombatants when conflict escalates. Brain imaging (functional magnetic resonance imaging) reveals that decreased harm avoidance is predicted by inhibition of the left middle frontal activity during selection of punishment decisions. Our findings provide insight into the cognitive and neural bases of decision-making involving civilian harm during intergroup conflict.

## Introduction

“Collateral damage” refers to civilian or noncombatant casualties and property damage due to aggressive actions during intergroup conflict such as a military operation during war^1^. Although conflicting with humanitarian interests and being prohibited by international laws^2^, “collateral damage” occurs massively in human societies^3, 4^ and raises serious moral and ethical concerns. For example, hundreds of civilians were killed and thousands of civilians were wounded due to NATO’s bombing intervention in the Kosovo conflict^5^. The ratio of civilian to military deaths during Iraq war is estimated to range between 3:1 and 8:1^4^. During the 2014 Israel-Gaza conflict over two thousand individuals were killed and 67% of the victims were civilians^6^. Given the substantial social consequences of civilian casualties, understanding its psychological constructs and brain underpinnings is of both theoretical and practical importance.

Civilian casualties are usually regarded as a side effect of aggressive acts following punishment decisions toward outgroup combatants during intergroup conflict. Punishment decision-making engages complicated cognitive and neural processes^7, 8^, including detection and generation of aversive experiences in the salience network (e.g., the anterior midcingulate and insula)^9^, integration of harm and intent into an assessment of blame in the theory-of-mind network (e.g., the medial prefrontal cortex and temporoparietal conjunction)^10^, and action selection/control in the central executive network (e.g., the lateral prefrontal cortex)^11^. In previous research, punishment decisions were made towards outgroup members indiscriminately as a whole in the context of intergroup conflict ^12–16^. This leaves an open issue related to civilian casualties, i.e., how to separate consequences of decision-making pertaining to outgroup combatants versus noncombatants so as to disentangle the cognitive/neural mechanisms underlying decision-making that causes harm to civilians. To address this issue would allow us to unravel psychological constructs characterizing the unique pattern of decision-making related to “collateral damage” during intergroup conflict. It is now known that a critical feature of real-world civilian casualties is that it increases when intergroup conflict escalates^17^. This suggests an approach to the understanding of cognitive and neural bases of punishment decision-making resulting in civilian casualties by examining psychological processes that vary along escalation of intergroup conflict.

Here, we propose two psychological constructs involved in punishment decision-making pertaining to outgroup members during intergroup conflict. Because the goal of combat during intergroup conflict is to punish or destroy outgroup combatants but to maintain minimal noncombatant casualties, as required by morality and international laws^2^, we hypothesize that harm preference and harm avoidance are two distinct psychological constructs that differentially characterize punishment decisions pertaining to outgroup combatants versus noncombatants during intergroup conflict (Hypothesis 1). Moreover, civilian casualties arising from intergroup conflict escalation may result from reduced harm avoidance pertaining to outgroup noncombatants (Hypothesis 2). That is, harm avoidance that inhibits harm to noncombatants during intergroup conflict is predicted to decrease when intergroup conflict escalates.

To test these hypotheses, we developed a behavioral paradigm in which participants were asked to make a decision by choosing electric shocks for punishing an outgroup member who was directly engaged in a conflict (an outgroup combatant) involving the distribution of resources. This punishment of the combatant was accompanied by varying degrees of harm to another outgroup member who was not involved in the conflict (an outgroup noncombatant). These manipulations in a laboratory setting simulate the nature of real-world civilian casualties during intergroup conflict in several aspects. First, the conflict between two groups was initiated by an unfair distribution of limited resources, which is consistent with research that shows that unfair money distributions induce conflict between two individuals or two groups^18^. Second, the participants had to make decisions to punish outgroup combatants, and these decisions also brought harm to outgroup noncombatants who were not involved in the conflict. Third, intergroup conflict was manipulated to vary across three (low, middle, or high) levels in our design. This design enables us not only to dissociate harm preference and harm avoidance by conducting statistical modeling of choices of electric shocks related to outgroup combatants and noncombatants, respectively, but also to estimate whether harm to outgroup noncombatants escalated when the level of intergroup conflict increased, similar to what happens during real-world intergroup conflicts^17^.

We examined how harm preference and harm avoidance varied along escalation of intergroup conflict in both minimal groups manipulated in a laboratory setting and two ethnic groups with conflicts in a real-world situation (i.e., Jewish and Palestinian groups during the Israeli-Palestinian conflict). Our statistical modeling of behavioral choices of punishment decisions showed consistent evidence for decreased harm avoidance pertaining to outgroup noncombatants during escalation of intergroup conflict. To further improve our understanding of neural underpinnings of punishment decision-making causing civilian casualties, we conducted an additional functional magnetic resonance imaging (fMRI) experiment to examine which neural network, e.g., the salience network, theory-of-mind network, or central executive network that contribute to punishment decision-making^7, 8^, is associated with harm preference or harm avoidance. In particular, we estimated neural underpinnings of punishment decision-making resulting in civilian casualties by examining brain activity associated with the decreased harm avoidance pertaining to outgroup noncombatants as intergroup conflict escalates.

## Results

### Decreased harm avoidance during escalation of intergroup conflict in a laboratory setting

To disentangle the psychological constructs related to civilian casualties from nonessential cognitive/affective processes engaged in punishment decision-making during intergroup conflict, in Experiment 1, we randomly assigned Chinese adults to a group conflict condition (N = 120) or an individual conflict (control) condition (N = 120). In the group conflict condition four participants of the same gender, who were strangers, were recruited simultaneously and tested in the same session. To build ingroup/outgroup relationships, the four participants were divided into two mini-groups (N = 2 in each mini-group) by wearing T-shirts of two different colors and playing a 1-hour card game (Phase 1 in Figure 1). This game required the participants to help ingroup members but to interrupt outgroup members so that one’s own group would reach a destination in advance (see Methods). After group formation in Phase 1, the participants completed a modified version of the Inclusion of Other in the Self Scale^19^ to assess their feelings of closeness between themselves and other participants. The same procedure was applied to the participants in the individual conflict condition except that there was no mini-group assignment and each participant competed against the other three in the game. Thus, no one shared group identity with others in the individual conflict condition, and the comparison of behavioral performance in group and individual conflict conditions allowed us to disentangle the effect specific to intergroup conflict in the group conflict condition by controlling other irrelevant factors. The analyses of the mean rating scores showed that the participants in the group conflict condition reported greater feelings of closeness to ingroup compared to outgroup members (mean ± s.d. = 4.92 ± 0.99 vs. 2.95 ± 1.14; F(1,116) = 232.40, *P* < 0.001, *η^2^_p_* = 0.667, 95% CI = 1.59, 2.61, BF_10_ > 150). The feelings of closeness to others in the individual conflict condition were lower than that pertaining to ingroup members in the group conflict condition (3.13 ± 1.03 vs. 4.92 ± 0.99; F(1,232) = 187.73, *P* < 0.001, *η^2^_p_* = 0.447, 95% CI = −2.05, −1.53, BF_10_ > 150) but did not differ significantly from that related to outgroup members in the group conflict condition (3.13 ± 1.03 vs. 2.95 ± 1.14; F(1,232) = 1.63, *P* = 0.203, *η^2^_p_* = 0.01, 95% CI = −0.10, 0.45, BF_01_ = 23.61). These results indicate that group formation increased the affective link between ingroup members rather than decreased the affective link to outgroup members relative to the control condition.

**Figure 1.**
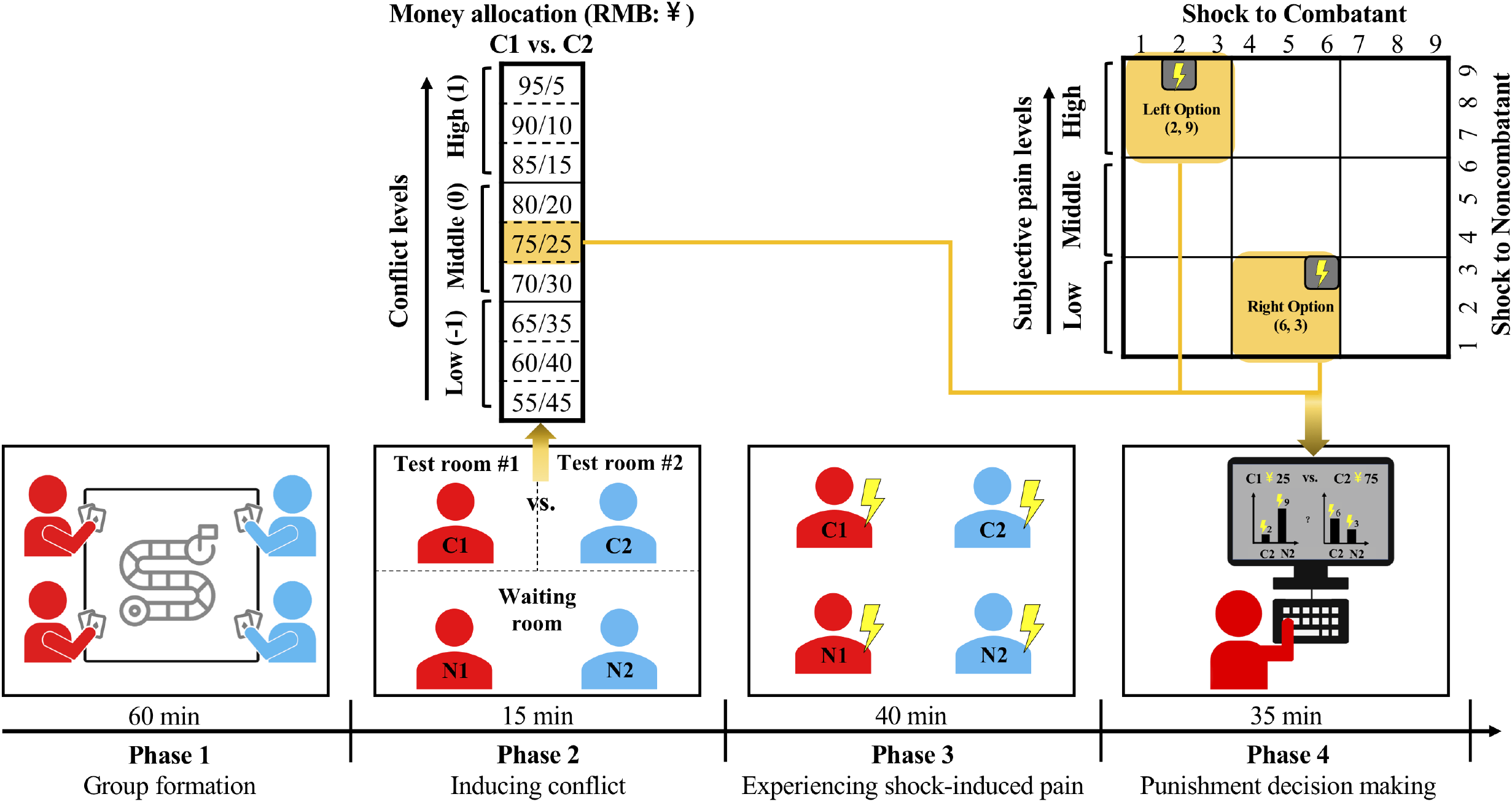
Illustration of the experimental procedure in Experiment 1. Group formation was manipulated in Phase 1 in which four participants were assigned to two groups to play a card game against an outgroup (the group conflict condition) or each participant played a card game against all the other three (the individual conflict condition). Group conflict was induced in Phase 2 in which C1 and C2 made decisions on the distribution of the bonus payment between C1 and C2. Three levels of conflict were defined based on unfairness of money distribution and were coded as low (−1), middle (0), and high (1) in data analyses. In Phase 3 the participants underwent a pain threshold test, respectively, and reported their subjective feelings of pain corresponding to different shock intensities. In Phase 4 each of the four participants was asked to make a punishment decision in a private room by choosing one from two options shown on the left and right side of a screen. Each option consisted of two electric shocks as punishment on the outgroup combatant and “collateral damage” to the outgroup noncombatants in the group conflict condition or punishment on a combatant and harm to a bystander in the individual conflict condition. Three levels of electric shocks were defined based on painful feelings corresponding to the stimuli (low, middle, high), resulting in nine combinations of shocks applied to combatant/noncombatant in the group conflict condition or combatant/bystander in the individual conflict condition.

Thereafter, in Phase 2 shown in Figure 1, one participant was randomly selected from each mini-group to play a combatant role in the group conflict condition (called C1 and C2, i.e. a combatant from group 1 and a combatant from group 2) and the remaining one in each group played the role of a noncombatant (called N1 and N2). In the individual conflict condition two participants were randomly selected as combatants (also called C1 and C2) with the left two being bystanders (also called N1 and N2). Thus C1 and N1 (or C2 and N2) shared group identity in the group conflict condition but not in the individual conflict condition. The following procedure was the same in the group and individual conflict conditions. Each noncombatant was informed of a 100 RMB payment for their participation. However, each combatant was told that, in addition to a basic payment of 50 RMB, he/she would share a 100 RMB bonus with the outgroup combatant as an additional payment and how much each person would obtain depends on the final distribution plan. C1 and C2 were then asked to make decisions privately on how to allot the bonus payment between C1 and C2 by choosing one of the distribution plans. Selection of a distribution plan of the bonus payment did not affect the payment for N1 and N2 so that N1 and N2 were not involved in the conflict and their noncombatant roles were not influenced by C1 and C2’s distribution plans. Selection of a distribution plan was repeated 108 times to make the participants believe various distribution plans used in the following punishment decision-making task. We defined three levels of conflict (low, middle, high) based on the distribution of the bonus payment between C1 and C2. In Phase 3 all participants went through a pain threshold test to rate their painful feelings (1 = slightly painful, 9 = extremely painful) corresponding to different intensities of electric shocks.

During the following punishment decision-making task (Phase 4 in Figure 1) each participant was tested in a private room to minimize social influences on their decision-making. Each trial began with the presentation of a plan of an unfair distribution (e.g., C1 and C2 receive 25 and 75 RMB, respectively), which was supposed to be given by an outgroup combatant (e.g., given by C2 in the case illustrated in Phase 4 in Figure 1) but was actually generated by a computer program. In the group conflict condition, the participant was instructed that, given the perceived distribution plan that was unfair to the ingroup combatant, they had to make a decision to punish the outgroup combatant. They were shown two punishment options, one on the left side and one on the right side of a screen, respectively (see illustration of Phase 4 in Figure 1). Each option consisted of two vertical bars indicating intensities (1 = slightly painful, 9 = extremely painful) of two electric shocks. One shock would be applied to the outgroup combatant as punishment and the other to the outgroup noncombatant as a collateral damage. There were three levels (i.e., 1-3, 4-6, or 7-9 corresponding to the low, middle, or high level of pain) of electric shocks. In either the left or right punishment option there were nine combinations of the two electric shocks applied to the outgroup combatant and noncombatant (i.e., 3 (low, middle, or high) levels of shocks to the combatant x 3 (low, middle, or high) levels of shocks to the noncombatant). These were the same in the individual conflict condition except that one shock would be applied to the combatant as punishment and the other to a bystander. The participants were told that the bonus and electric shock in one of their decisions would be randomly selected to apply to the target combatant and noncombatant after the experiment.

The distribution plan and the options of electric shocks were manipulated to balance conflict levels and punishment options. Each participant completed 108 trials of punishment decision-making with 3 levels of conflict based on the given distribution plan (i.e., leaving 55-65, 70-80, or 85-95 RMB for oneself, corresponding to low, middle, or high conflict) and three (low, middle, or high) levels of electric shocks. The intensities of the two electric shocks applied to outgroup combatants and noncombatants in the group conflict condition (or to combatants and bystanders in the individual conflict condition) varied independently across trials. There were 108 different combinations of conflict level and electric shock intensity that were presented to each participant in a random order.

Participants’ choices of electric shocks were subject to logistic regression analyses^20^. Because we hypothesized that outgroup combatants and noncombatants are treated differently in terms of harm preference and harm avoidance during intergroup conflict, we included four independent variables in the regression model to account for the probability that participants would select the left option (*p*) or the probability that participants would select the right option (1*-p*) in each trial (see Method, Supplementary Fig. 1 and Supplementary Table 1) for the methods and results of model comparisons). These included the difference in intensities of electric shocks between left and right options applied to the combatant (ΔS_c_) and the noncombatant (ΔS_n_), the interactions between ΔS_c_ and level of conflict (L) and between ΔS_n_ and L. The logistic regression model was defined as following:

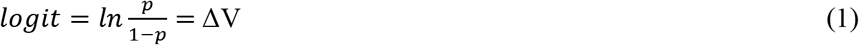

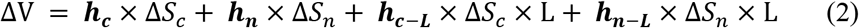

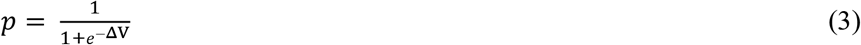

A larger *p* is associated with higher possibility of selection of the left option.

The regression coefficients of these independent variables in this logistic regression model are associated with different psychological constructs. A positive ***h_c_*** or ***h_n_*** value indicates harm preference — a propensity to select a stronger electric shock from two options to punish a target. A negative ***h_c_*** or ***h_n_*** value indicates harm avoidance — a propensity to select a weaker electric shock from two options to punish a target. ***h_c-L_*** (or ***h_n-L_***) manifests how ***h_c_*** (or ***h_n_***) varies along escalated levels of conflict and positive (or negative) ***h_c-L_*** or ***h_n-L_*** values indicate increase (or decrease) of ***h_c_*** or ***h_n_***. To reduce potential biases in estimation and to allow for finite parameter estimates for data with separation, we performed logistic function estimation using Firth’s penalized-likelihood logistic regression (*logistf* function in R)^21, 22^. The logistic regression model was estimated and the four regression coefficients were extracted for each participant for the second group-level t-tests. We also conducted Bayesian factor analyses^23^ to assess the likelihood of data driven alternative hypothesis over the likelihood of data driven null hypothesis (BF_10_) or the reverse (BF_01_) (see Methods for details).

If harm preference and harm avoidance characterize punishment decisions pertaining to outgroup combatants and noncombatants, respectively, we would expect a positive value of ***h_c_*** (harm preference to combatants) but a negative value of ***h_n_*** (harm avoidance related to noncombatants). In addition, if collateral damage during escalation of intergroup conflict arises from reduced harm avoidance pertaining to outgroup noncombatants, we predicted a positive value of ***h_n-L_***.

As expected, the results of one sample t-tests (all two-tailed) showed that ***h_c_*** was significantly larger than zero in both the group conflict condition (mean ± s.d. = 0.18 ± 0.80, t(119) = 2.47, *P* = 0.015, FDR corrected, Cohen’s *d* = 0.23, 95% CI = 0.04, 0.32; BF_10_ = 1.84, only FDR corrected results were reported in the following) and the individual conflict condition (0.31±0.68, t(119) = 4.95, *P* < 0.001, Cohen's d = 0.45, 95% CI = 0.19, 0.43; BF10 > 150, Figure 2, Supplementary Fig. 2), but did not differ significantly between the two conditions (two sample test: t(238) = −1.35, *P* = 0.358, Cohen's *d* = 0.17, 95% CI = −0.32, 0.06; BF_01_ = 3.00, see Figure 2). However, ***h_n_*** was significantly smaller than zero in both the group conflict condition (−0.52±0.53, t(119) = −10.71, *P* < 0.001, Cohen’s *d* = 0.98, 95% CI = −0.61, −0.42; BF_10_ > 150) and in the individual conflict condition (−0.55±0.50, t(119) = −12.14, *P* < 0.001, Cohen’s *d* = 1.11, 95% CI = −0.64, −0.46; BF_10_ > 150), but did not differ significantly between the two conditions (t(238) = 0.55, *P* = 0.584, Cohen's *d* = 0.07, 95% CI = −0.09, 0.17; BF_01_ = 6.14). These results provide evidence for harm preference for combatants (indicated by a positive ***h_c_***) but harm avoidance related to noncombatants (indicated by a negative ***h_n_***) on average across the three conflict levels.

**Figure 2.**
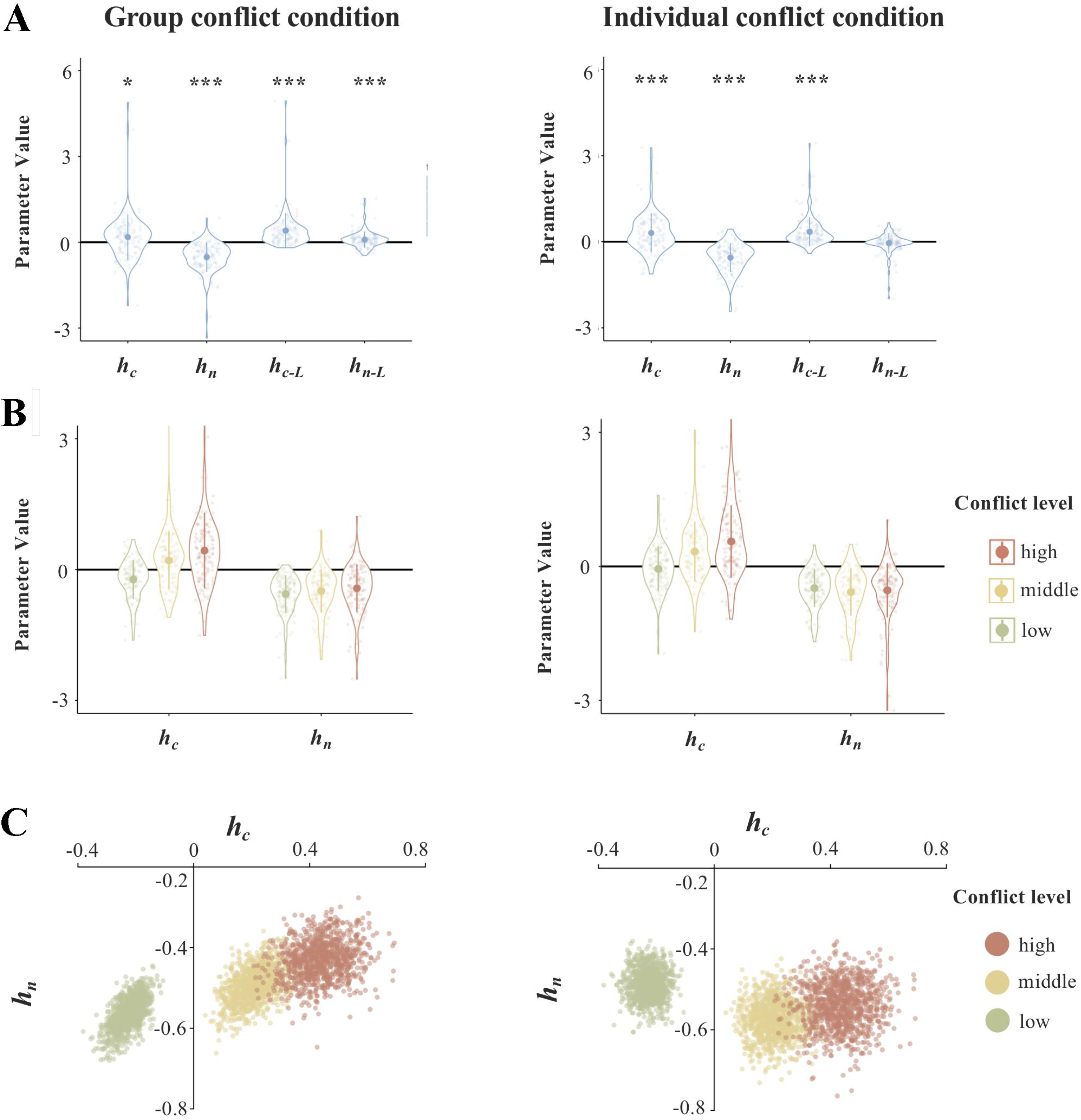
Results of logistic regression parameter analyses in the group and individual conflict conditions in Experiment 1. (**A**) Violin plots show means (big dots), s.d. (bars), and distributions of parameter values. * *P* < 0.05; *** *P* < 0.001, FDR corrected. Each small dot represents an individual participant. (**B**) Bar charts illustrate variations of ***h_c_*** and ***h_n_*** that were calculated at each level of conflict and in different conditions. ***h_n_*** was significantly smaller than zero in both the group conflict condition (−0.52±0.53, t(119) = −10.71, *P* < 0.001, Cohen’s *d* = 0.98, 95% CI = −0.61, −0.42; BF_10_ > 150) and individual conflict condition (−0.55±0.50, t(119) = −12.14, *P* < 0.001, Cohen’s *d* = 1.11, 95% CI = −0.64, −0.46; BF_10_ > 150). ***h_n-L_*** was significantly larger than zero in the group conflict condition (0.09±0.29, t(119) = 3.56, *P* < 0.001, Cohen’s *d* = 0.33, 95% CI = 0.04,0.15; BF_10_ = 36.87) but not in the individual conflict condition (−0.05±0.35, t(119) = −1.50, *P* = 0.135, Cohen's *d* = 0.14, 95% CI = −0.11, 0.02; BF_01_ = 3.30). (**C**) The results of bootstrapping analyses illustrate variations of sample distributions of ***h_c_*** and ***h_n_*** along three levels of conflict in the group and individual conflict conditions. We conducted repeated random resampling with replacement from the available data and the mean of this bootstrapped sample was then calculated and plotted in a two-dimensional space defined by ***h_c_*** and ***h_n_***. The procedure was replicated for 1000 times to generate an empirical approximation of the sampling distribution. ***h_c_***, ***h_n_***, ***h_c-L_*** and ***h_n-L_*** are regression coefficients in the logistic regression model. A positive ***h_c_*** or ***h_n_*** value indicates harm preference whereas a negative ***h_c_*** or ***h_n_*** value indicates harm avoidance. ***h_c-L_*** (or ***h_n-L_***) manifests how ***h_c_*** (or ***h_n_***) varies along escalated levels of conflict. Positive (or negative) ***h_c-L_*** or ***h_n-L_*** values indicate increase (or decrease) of ***h_c_*** or ***h_n_***.

The analyses of ***h_c-L_*** showed similar patterns in the group and individual conflict conditions whereas the analyses of ***h_n-L_*** revealed different patterns between the two conditions. ***h_c-L_*** was significantly larger than zero in both the group conflict condition (0.41±0.61, t(119) = 7.33, *P* < 0.001, Cohen’s *d* =0.67, 95% CI = 0.30, 0.52; BF_10_ > 150) and individual conflict condition (0.35±0.51, t(119) = 7.58, *P* < 0.001, Cohen’s *d* =0.69, 95% CI = 0.26, 0.44; BF_10_ > 150), but did not differ significantly between the two conditions (t(238) = 0.79, *P* = 0.576, Cohen's *d* =0.10, 95% CI = −0.09, 0.20; BF_01_ = 5.28). The results suggest that, for both group conflict and individual conflict conditions, harm preference (indicated by ***h_c_***) towards combatants increased when conflict escalated. Most interestingly, ***h_n-L_*** was significantly larger than zero in the group conflict condition (0.09±0.29, t(119) = 3.56, *P* < 0.001, Cohen's *d* = 0.33, 95% CI = 0.04,0.15; BF_10_ = 36.87) but not in the individual conflict condition (−0.05±0.35, t(119) = −1.50, *P* = 0.135, Cohen's *d* = 0.14, 95% CI = −0.11, 0.02; BF_01_ = 3.30). A two-sample t-test further confirmed the significant difference in ***h_n-L_*** between the group and individual conflict conditions (t(238) = 3.42, *P* = 0.003, Cohen’s *d* = 0.44, 95% CI = 0.06, 0.22; BF_10_ = 32.28). These results provide evidence that harm avoidance toward noncombatants decreased (i.e., ***h_n_*** (harm avoidance related to noncombatants) shifted toward less negative) along escalated conflict in the group conflict condition but not in the individual conflict condition. This finding supports the idea that decreased harm avoidance functions as a psychological construct related to civilian casualties that occurs specifically during intergroup conflict.

To test how well the logistic regression model could predict individuals’ punishment decisions, we compared participants’ choices and the model prediction of a choice in each trial (e.g., probability to choose left or right option with 0.5 as the threshold). The results showed that the logistic regression model correctly predicted 85.03 ± 7.77% of choices, which is significantly higher than the chance level (t(239) = 69.85, *P* < 0.001, Cohen’s *d* = 4.51, 95% CI = 0.84, 0.86; BF_10_ > 150, see Supplementary Fig. 1). The results indicate that the logistic regression model (Model 1) fits well with the participants' actual performances during decision making.

### Decreased harm avoidance during escalation of intergroup conflict in a real-world context

The results of Experiment 1 suggest decreased harm avoidance toward outgroup noncombatants during intergroup conflict escalation in a laboratory setting. In Experiment 2 we examined whether this account applies to collateral damage during intergroup conflict in a real-world context by testing Jewish (N = 106) and Palestinian participants (N = 106) in Israel. The Israeli-Palestinian conflict is one of the most challenging intergroup conflicts in the world and threatens numerous civilians’ lives^6,24–26^. The study of participants from these two groups allowed us to further understand the role of decreased harm avoidance in civilian casualties in a real-world intergroup conflict. Experiment 2 was carried out during the year 2018 when repeated clashes with numerous casualties occurred between the two sides. Because of the difference in national and religious identity between the two ethnic groups, we divided 2 Jewish participants and 2 Palestinian participants (all were of the same gender and strangers to each other) into two mini-groups on each testing day. The participants were blind to what the study was about during the recruitment. To highlight participants' ethnic identity, after introducing one's own ethnicity, all participants were presented with images of items or celebrities/holy places that remind them of the ethnicity-specific cultural elements^27^. The participants from each mini-group were asked to jointly observe each picture and rate their feelings of picture-induced belonging and excitement (1 = very little, 9 = very much). The rating scores were high for both ethnic groups and higher in Palestinian participants than in Jewish participants (see Supplementary information, Results 1 and 2, Supplementary Tables 2 and 3, and Supplementary Fig. 3).

Thereafter, participants completed the money distribution task, pain threshold test, and punishment decision-making task. The procedures of these tasks were the same as those in Experiment 1 except that the participants were paid with a corresponding amount of Shekels. Similarly, a logistic regression model was constructed to estimate each participant's performance during punishment decision-making. We first assessed harm preference and harm avoidance characterizing the whole sample by pooling together the two ethnic groups’ data and then examined possible group differences. One sample t-tests conducted on the whole sample showed that ***h_c_*** (harm preference to combatants) did not differ significantly from zero (mean ± s.d. = −0.07±0.55, t(211) = −1.76, *P* = 0.081, Cohen’s *d* = 0.12, 95% CI = −0.14, 0.01; BF_01_ = 2.88), whereas ***h_n_*** (harm avoidance related to noncombatants) was significantly smaller than zero (−0.50±0.46, t(211) = −15.67, *P* < 0.001, Cohen's *d* = 1.08, 95% CI = −0.56, −0.44; BF_10_ > 150, Figure 3, Supplementary Fig. 4 and 5). Moreover, these effects were significantly smaller in Jewish than in Palestinian participants (***h_c_***: −0.19±0.55 vs. 0.06±0.51, t(210) = −3.49, *P* = 0.001, Cohen’s *d* = 0.57, 95% CI = −0.38, −0.14; BF_10_ > 150; ***h_n_***: −0.63±0.49 vs. −0.37±0.39, t(210) = −4.17, *P* < 0.001, Cohen's *d* = 0.57, 95% CI = −0.38, −0.14; BF_10_ > 150). The results of one sample t-tests conducted on the whole sample revealed that both ***h_c-L_*** and ***h_n-L_*** were significantly larger than zero (***h_c-L_***: 0.19±0.28, t(211) = 9.84, *P* < 0.001, Cohen's *d* = 0.68, 95% CI = 0.15, 0.23; BF_10_ > 150; ***h_n-L_***: 0.06±0.28, t(211) = 3.04, *P* = 0.003, Cohen's *d* = 0.21, 95% CI = 0.02, 0.10); BF_10_ = 6.66). These effects did not differ significantly between Jewish and Palestinian participants (***h_c-L_***: 0.19±0.30 vs. 0.19±0.26, t(210) = 0.14, *P* = 0.888, Cohen’s *d* = 0.02, 95% CI = −0.07, 0.08; BF_01_ = 6.62; ***h_n-L_***: 0.05±0.34 vs. 0.07±0.20, t(210) = −0.57, *P* = 0.756, Cohen's *d* = 0.08, 95% CI = −0.10, 0.05; BF_01_ = 5.72).

**Figure 3.**
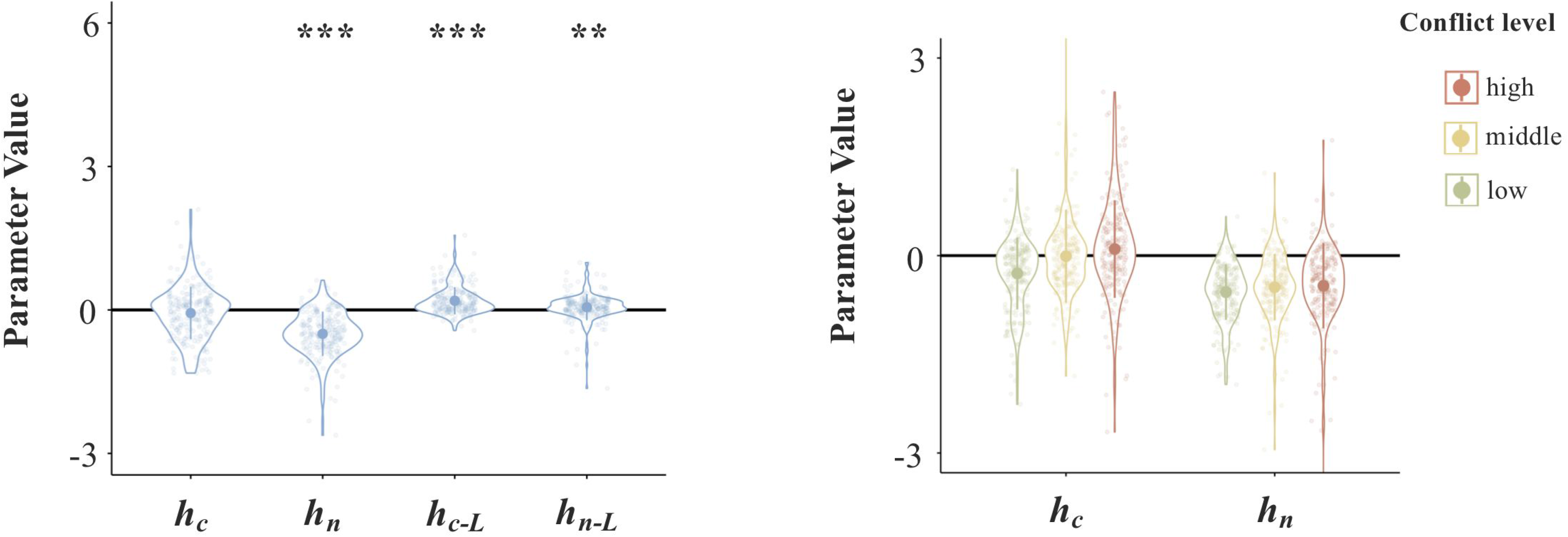
Results of logistic regression parameter analyses of the whole sample in Experiment 2. The left panel shows violin plots of mean (big dots), s.d. (bars), and distributions of parameter values. ** *P* < 0.005; *** *P* < 0.001, FDR corrected. Each small dot represents an individual participant. The right panel illustrates variations of ***h_c_*** and ***h_n_*** along increased conflict levels. ***h_n_*** was significantly smaller than zero (−0.50±0.46, t(211) = −15.67, *P* < 0.001, Cohen's *d* = 1.08, 95% CI = −0.56, −0.44; BF_10_ > 150) and ***h_n-L_*** was significantly larger than zero (0.06±0.28, t(211) = 3.04, *P* = 0.003, Cohen’s *d* = 0.21, 95% CI = 0.02, 0.10); BF_10_ = 6.66).

Similarly, the logistic regression model predicted a high proportion of punishment decisions across trials that was significantly higher than the chance level (83.89%±8.39%, t(211) = 58.79, *P* < 0.001, Cohen's *d* = 4.04, 95% CI = 82.75%, 85.02%; BF_10_ > 150, Supplementary Fig. 1), indicating that the logistic regression model fits well with actual performance in the sample tested in a real-world situation.

Together, the results in Experiments 1 and 2 provide evidence for decreased harm avoidance toward noncombatants when intergroup conflicts escalate in both a laboratory setting and a real social context. Moreover, the same logistic regression model fits the empirical data of punishment decision-making during intergroup conflict in both experiments. These results suggest that decreased harm avoidance provides a common cognitive basis of decision-making involving civilian casualties in a laboratory setting and a real-world social context. Notably, our participants in a laboratory setting but not in a real-world situation showed harm preference and Jewish participants even exhibited harm avoidance toward outgroup combatants (see Supplementary Fig. 4). Punishment decisions seemed to be made more carefully in a real social context of intergroup conflict than in a laboratory setting possibly due to higher stakes in the latter situation. It is also possible that the procedure used in our study created demand characteristics that resulted in social desirability pressures to appear moral. Future research should examine these distinct possibilities and replicate them with other samples embedded in real world conflicts.

### Neural correlates of decreased harm avoidance during intergroup conflict escalation

In Experiment 3, we further investigated neural correlates of the decreased harm avoidance during intergroup conflict by testing an independent sample of Chinese adults. The experimental procedures were the same as those in Experiment 1 except that participants were scanned using fMRI during the punishment decision task and the amounts of the basic payment and additional bonus payment were doubled. Participants were randomly assigned to the group conflict condition (N = 64) and the individual conflict condition (N = 64). Logistic regression analyses of the behavioral data replicated the findings of decreased harm avoidance in punishment decision-making related to outgroup noncombatants during conflict escalation in the group conflict condition (Figure 4A, see Supplementary information, Result 3, Supplementary Table 4 and Supplementary Fig. 6 for details).

**Figure 4.**
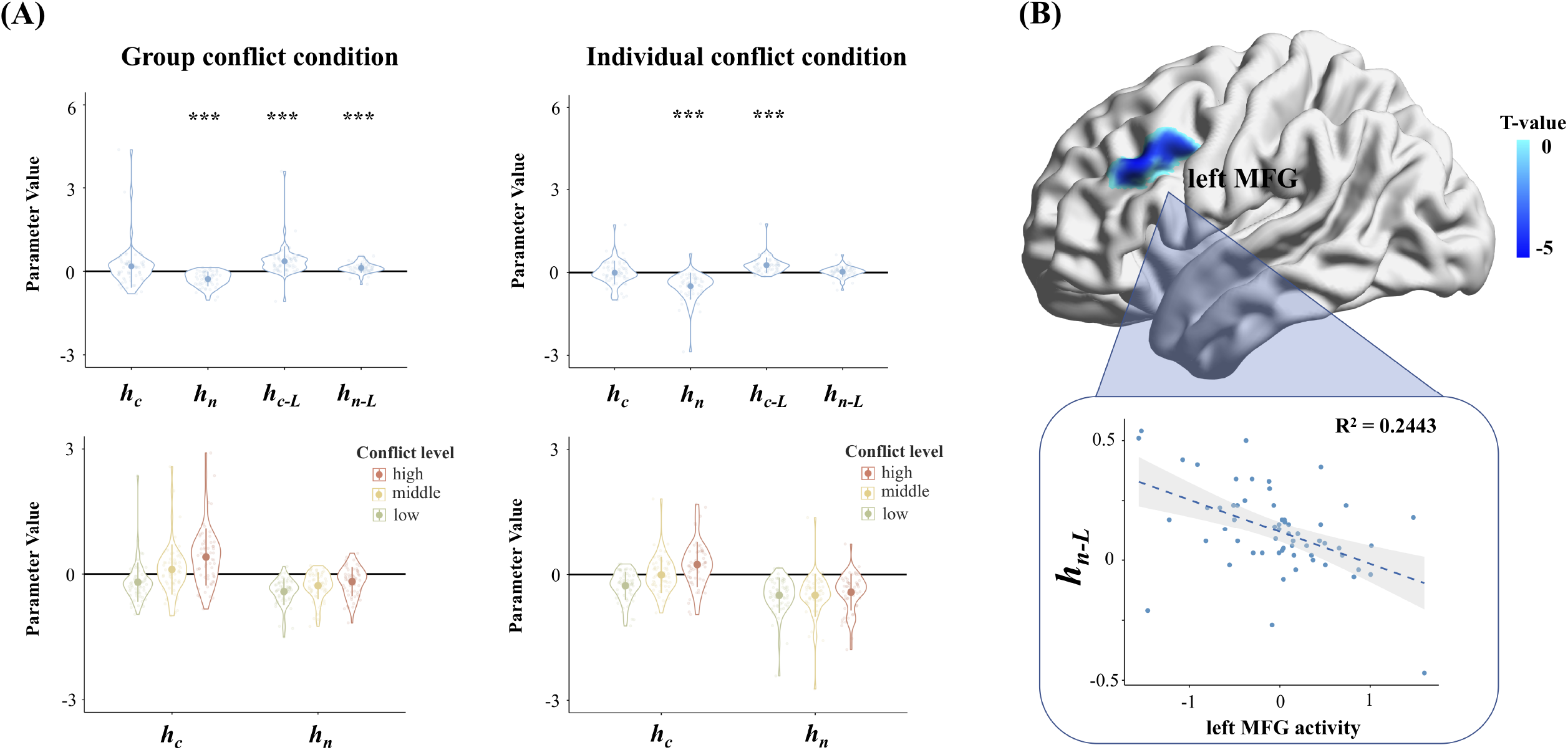
Behavioral and fMRI results in Experiment 3. (**A**) The results of logistic regression parameter analyses in the group and individual conflict conditions, respectively. The top panels show violin plots of mean (big dots), s.d. (bars), and distributions of parameter values. *** *P* < 0.001, FDR corrected. Each small dot represents an individual participant. The bottom panels illustrate variations of ***h_c_*** and ***h_n_*** along increased conflict levels. ***h_n_*** was significantly smaller than zero in both the group conflict condition (−0.28±0.27, t(59) = −8.04, *P* < 0.001, Cohen's *d* = 1.04, 95% CI = −0.35, −0.21; BF_10_ > 150) and individual conflict condition (−0.50±0.49, t(58) = −7.83, P < 0.001, Cohen's d = 1.02, 95% CI = −0.62, −0.37; BF_10_ > 150). ***h_n-L_*** was significantly larger than zero in the group conflict condition (0.12±0.18, t(59) = 5.36, *P* < 0.001, Cohen's *d* = 0.69, 95% CI = 0.08,0.17; BF_10_ = 36.87) but not in the individual conflict condition (0.02±0.19, t(58) = 0.94, *P* = 0.467, Cohen's *d* = 0.12, 95% CI = −0.03,0.07; BF_01_ = 0.22). (**B**) The decreased left MFG activity during selection of punishment decisions predicted larger ***h_n-L_*** in the group conflict condition (a voxel-level threshold *P* < 0.001, cluster-level threshold *P* < 0.05, FWE corrected). The top panel illustrates the location of the left MFG. The bottom panel illustrates variation of the left MFG activity across individuals’ ***h_n-L_***.

The first-level whole-brain general linear model (GLM) analyses of fMRI data estimated brain activities associated with the parametric modulators including ΔS_c_, ΔS_n_, ΔS_c_ * L, and ΔS_n_ * L in each participant. We conducted four second-level whole-brain regression analyses to test brain regions associated with individual differences in parameters estimated using the logistic regression modelling. The second-level regression analyses included the contrast images of parametric modulation effects of ΔS_c_ (with the covariate ***h_c_***), ΔS_n_ (with the covariate ***h_n_***), ΔS_c_ * L (with the covariate ***h_c-L_***), and ΔS_n_ * L (with the covariate ***h_n-L_***) of each participant. Most interestingly, the second-level analyses showed that a larger ***h_n-L_*** was significantly correlated with decreased activity in the left middle frontal gyrus (MFG, MNI coordinates x/y/z = −48/28/32, a voxel-level threshold *P* < 0.001, cluster-level threshold *P* < 0.05, FWE corrected, Figure 4B) in the group conflict condition but not in the individual conflict condition, suggesting a functional role of the left MFG in controlling harm avoidance pertaining to outgroup noncombatants when intergroup conflict escalated.

To further confirm that the difference in the association between the left MFG activity and ***h_n-L_*** was statistically significant when directly comparing the group and individual conflict conditions, we conducted a moderation analysis of the association between ***h_n-L_*** and the left MFG activity. This analysis searched for regions in the left frontal lobe defined by a mask in the Brainnetome Atlas^28^ in which the association between ***h_n-L_*** and the left MFG activity differed significantly between the group and individual conflict conditions. The result revealed a significant activation in a region in which the voxel with peak activation was located at −48/28/30 (z = 3.65, voxel-level threshold *P* < 0.05 with small volume FWE correction). This region overlaps with the left MFG shown in the second-level whole-brain regression analyses and provides evidence that intergroup conflict significantly enhanced the coupling between ***h_n-L_*** and the left MFG activity compared to the control condition.

The second-level whole-brain regression analysis across the participants using the contrast image of parametric modulation effects of ΔS_n_ and ***h_n_*** showed that ***h_n_*** (harm avoidance related to noncombatants) was associated with increasing activity in the left precentral cortex in the group conflict condition (−36/-30/56) and in the bilateral precentral cortices in the individual conflict condition (−36/-14/64 and 60/8/32, a voxel-level threshold *P* < 0.001, cluster-level threshold *P* < 0.05, FWE corrected, Figure 5A), but these activations did not differ significantly between the group and individual conflict conditions. Because there was no significant difference in ***h_c_*** (harm preference to combatants) and ***h_c-L_*** between the group and individual conflict conditions, we pooled fMRI data in the two conditions to examine neural correlates of ***h_c_*** and ***h_c-L_***, respectively. The results of the second-level whole-brain regression analyses showed that a larger ***h_c_*** was associated with increased activity in the left temporoparietal junction (−50/−40/26), left prefrontal cortex (−28/36/32), midcingulate (14/2/48), and bilateral occipital cortex (−34/−76/26 and 34/−74/16), whereas increased activity linked to ***h_c-L_*** was observed only in the right occipital cortex (24/−54/−10, voxel-level threshold *P* < 0.001, cluster-level threshold *P* < 0.05, FWE corrected, Figure 5B). Apparently, these activations do not overlap with the left MFG which seems to be specifically associated with decreased harm avoidance related to outgroup noncombatants during intergroup conflict. Taken together, these neuroimaging results uncovered brain activity that was specifically associated with collateral damage during intergroup conflict (i.e., the left MFG activity). Our neuroimaging results also revealed brain activities underlying punishment decision making that, however, were similarly observed in the group and individuals conflict conditions.

**Figure 5.**
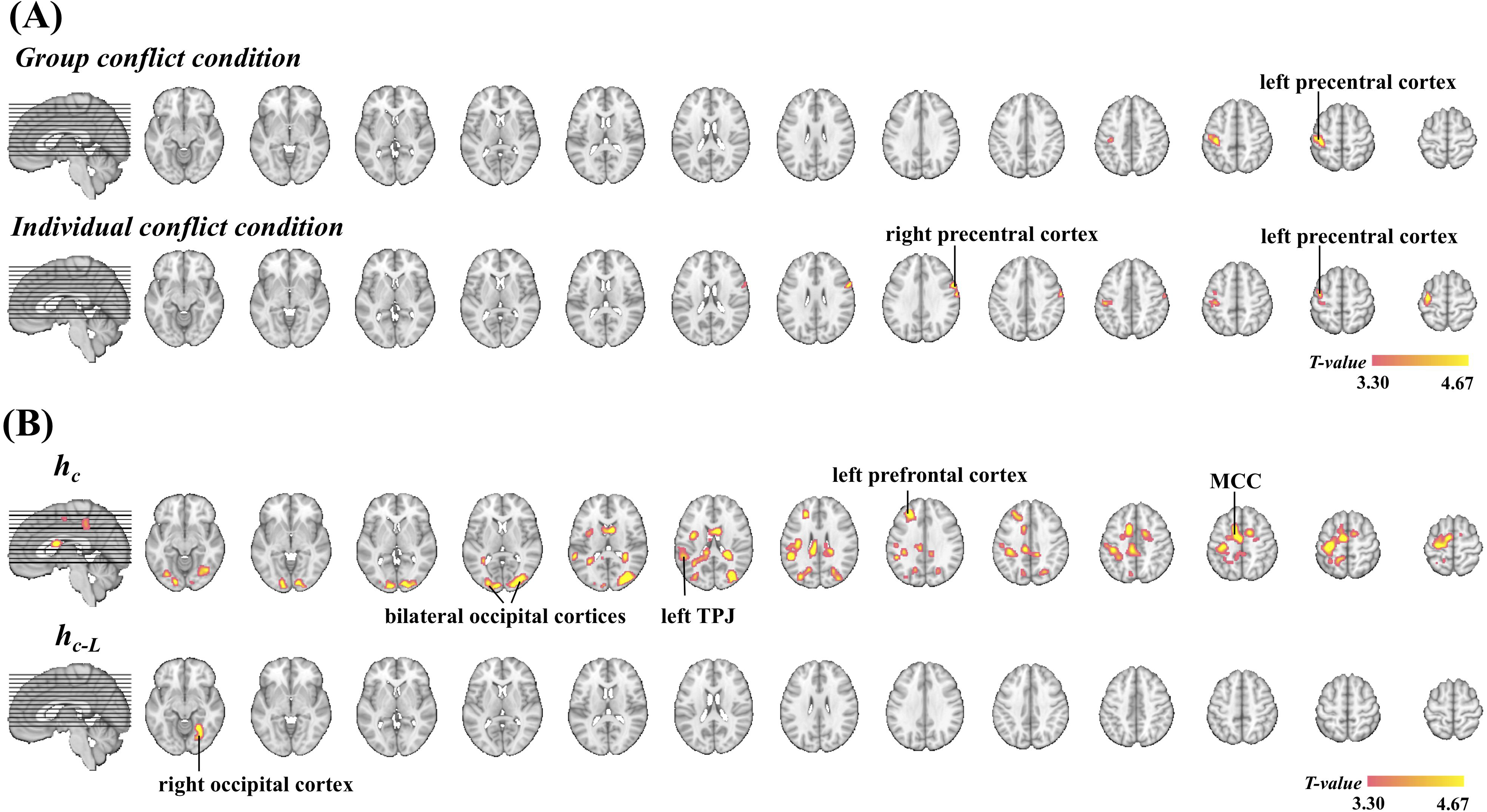
fMRI results of Experiment 3. (A) Neural correlates of ***h_n_*** in the group and individual conflict groups, respectively. (B) Neural correlates of ***h_c_*** and ***h_c-L_*** when combining group and individual conflict groups. Brain activations were defined by combining a whole-brain voxel-level threshold of *P* < 0.001 and a cluster-level threshold at *P* < 0.05 FWE corrected. MCC=midcingulate cortex; TPJ=temporoparietal junction.

## Discussion

The present study combined functional brain imaging and statistical modeling of punishment decision-making to examine the cognitive and neural bases of decision-making resulting in civilian casualties during intergroup conflict. Unlike previous research of intergroup conflict that measured punishment decisions upon outgroup members as a whole^12–16^, our design distinguished between outgroup combatants and noncombatants in punishment decision making. By comparing individuals’ behavioral performance in group-conflict and individual-conflict conditions, we highlighted a psychological construct, i.e., decreased harm avoidance, which was specifically related to selection of punishment decisions that led to civilian casualties when intergroup conflict escalated. Our fMRI results further uncovered the neural correlate of this psychological construct in the left MFG.

Our statistical modeling revealed a significantly negative value of the parameter ***h_n_*** as an index of harm avoidance during the selection of electric shocks that were supposed to be possibly applied to noncombatants. This finding indicates that people tended to give weaker electric shocks to or to avoid harming others who were not directly engaged in aggressive actions during conflict. This was true during conflicts between two groups or between two individuals and is consistent with moral imperatives and international laws in human societies^29, 30^. Nevertheless, the statistical modeling revealed a positive value of the parameter ***h_n-L_*** in the situation of intergroup conflict across three experiments. The positive ***h_n-L_*** suggests that harm avoidance that inhibits harm to noncombatants during intergroup conflict tends to decrease when intergroup conflict escalates.

It has been speculated that, as conflict escalates, decision makers are less likely to differentiate between combatants and noncombatants due to depleted cognitive resources and have to deal with the incidental cognitive dissonance^17^. These hypothesized cognitive mechanisms underlying civilian casualties, however, have not been tested empirically. Our empirical findings uncover a psychological construct, i.e., harm avoidance, which varies across levels of intergroup conflict in a fashion similar to actual civilian casualties. Furthermore, we showed that decreased harm avoidance during conflict escalation was evident in individuals facing conflict between an ingroup member and an outgroup member but not in those facing conflicts between two individuals who had no ingroup/outgroup relationships with the decision maker. This provides further evidence that, rather than reflecting changes of cognitive processes that are general to any conflict, the decreased harm avoidance specifically characterizes punishment decision-making that leads to civilian casualties during escalation of intergroup conflict. In addition, our results demonstrated a positive value of ***h_n-L_*** in both a laboratory setting in China and in a natural conflict context in Israel. Thus a similar psychological construct may underlie civilian casualties during escalation of intergroup conflict in both a laboratory setting and a real-world context.

Our statistical modeling also revealed a significantly positive value of the parameter ***h_c_*** as an index of harm preference for outgroup combatants during selection of electric shocks in a laboratory setting but not in a real social context of intergroup conflict. However, the statistical modeling showed a significant positive value of the parameter ***h_c-L_*** across the three experiments in both the group and individual conflict conditions. The positive ***h_c-L_*** suggests that our participants tended to choose stronger electric shocks to punish an individual if this person had proposed a more unfair money distribution plan. The increased harm preference as a function of conflict escalation might have different psychological causes in the group and individual conflict conditions. In the group conflict condition, when facing a more unfair money distribution plan, noncombatants might choose stronger electric shocks to protect ingroup members’ benefit, similar to previous findings of ingroup favoritism in decision-making^31–33^. In the individual conflict condition, however, unfair money distribution between two individuals did not directly affect a bystander’s payoff and punishment decision-making was possibly made to protect social norms, similar to third-party punishment^34^. Our data analyses did not show consistent evidence for differences in parameter estimates of harm preference and harm avoidance between combatants and noncombatants as decision makers. It is likely that the motives to help ingroup and to protect social norms dominated punishment decision-making whereas one’s own payment was much less taken into consideration in our participants. Future research requires a new behavioral paradigm that helps to reveal how combatants and noncombatants make punishment decisions differently.

We integrated fMRI and statistical modeling of punishment decision-making to examine neural correlates of harm avoidance and harm preference during intergroup conflict. Our fMRI results showed that ***h_c_*** and ***h_n_*** were linked to distinct neural circuits, i.e., ***h_n_*** was associated with increasing activity in the precentral cortex whereas ***h_c_*** was associated with increased activity in the left temporoparietal junction, left prefrontal cortex, and midcingulate. These results provide further neuroimaging evidence that harm avoidance and harm preference represent two distinct psychological processes supported by different neural underpinnings. Most importantly, we showed that the decreased left MFG activity predicted a larger ***h_n-L_*** and this effect was specific to the participants engaged in intergroup conflict. This finding suggests a key functional role of the left MFG in controlling harm avoidance related to civilian casualties during intergroup conflict. Specifically, inhibition of the left MFG might lead to failure to avoid harm to noncombatants when making punishment decisions during high intergroup conflict. The functional role of the left MFG during intergroup conflict is consistent with previous fMRI findings. For example, the left MFG activity increases with the complexity of different cognitive tasks and lesions of this brain region result in performance deficits in different cognitive tasks^35^. The lateral frontal cortex also represents the value of employing cognitive control and plays an important role in selection/control of a specific punishment decision^8, 11^ and in value-based learning and decision-making^36^. These findings agree upon a general function of the left MFG in executive control of decision-making and behavioral responses that is independent of task domains.

Our fMRI results revealed increasing activation in the precentral cortex as neural correlates of harm avoidance related to noncombatants as indexed by ***h_n_***. This is consistent with the functional role of this brain region in general behavioral inhibition^37^. Harm preference toward combatants as indexed by ***h_c_*** was associated with increased activity in the left temporoparietal junction, left prefrontal cortex, and midcingulate. These brain regions are involved in punishment decision-making to detect aversive experiences and to infer others’ mental states^7–10^. Similar mental processes may support harm preference regardless of when conflicts occur between two groups or between two individuals. However, although the motives to punish outgroup combatants and to avoid harming to noncombatants may coexist, inhibition of the left MFG activity is specifically related to the decreased harm avoidance as a psychological construct underlying civilian casualties when intergroup conflict escalates.

We also considered revenge as a potential motivator of punishment in this study. Revenge is usually manifested in taking actions of harming someone in retaliation for an injury^38, 39^. Our recent fMRI research showed that the medial prefrontal activity in response to ingroup pain caused by outgroup members predicted the propensity to give painful electric shocks to outgroup members^40^. In the current study the participants made choices by considering shocks to both the outgroup combatant and noncombatant. While our statistical modeling of punishment decision-making suggested harm preference toward the outgroup combatant, the results also suggest harm avoidance toward the outgroup noncombatant. It is thus unlikely that the participants made their choices only by considering retaliation upon the outgroup combatant. Moreover, revenge cannot explain why harm avoidance related to outgroup noncombatants decreased when intergroup conflict escalated since the outgroup noncombatants did not bring any harm to the decision maker in any case. We also assessed the possibility that the participants made their choices by considering retaliation upon the outgroup combatant and noncombatant as a whole (or group revenge) by constructing a different model (i.e., Model 3 reported in Supplementary Fig. 1 and Supplementary Table 1). In this model the sum of shock intensities to the combatant and the noncombatant (i.e., group revenge) was used in a regression model to predict the participants choices. The results suggest that this ‘group revenge’ model is not as good as the model reported in the main text in predicting the participants’ decisions. It appears that revenge cannot fully explain punishment decision making in a context when a punishment decision is related to both outgroup combatants and noncombatants during intergroup conflict.

Previous research on mechanisms of intergroup conflict usually took all outgroup members as one unit. Perceiving the outgroup as a competitor for limited resources^41^ or a threat to the ingroup^42^ may trigger intergroup conflict. Dehumanization of outgroup members^43^ may further drive discrimination and conflict. Humans may evolve ingroup favoritism in implicit attitudes^44^ and empathy^45–50^ and aggressiveness against outgroups as genetic or cultural traits^51^. While these traits help individuals to survive, humans as a rational species are confronted with the contradiction between harming outgroup combatants as enemies and protecting outgroup noncombatants as innocents during intergroup conflict. Our findings provide a neurocognitive account of decision-making that underlies civilian casualties during intergroup conflict.

While our findings contribute to the understanding of neurocognitive underpinnings of decisions involving harm to noncombatants in real-world situations, several limitations of our findings should be acknowledged. First, civilian casualties in real-world situations usually arise from armed attacks toward outgroup combatants and such attacks cost lives of and bring long-term physical/mental harms to outgroup combatants and noncombatants. The harm that participants intended to apply to the outgroup in our paradigm, however, was much weaker than those in real-world situations due to ethical requirements. Nevertheless, despite the differences in degree of harm, our paradigm captures the key features of collateral damages in real-world situations (i.e., increasing along conflict escalation). Besides, previous research has shown that conflict expenditures of groups can be substantially larger than those of conflict between individuals and allowing group members to punish each other results in even larger conflict expenditures^52^. In our paradigm, however, conflict expenditures (distributed money between the two combatants) were controlled in the group and individual conflict conditions as an irrelevant variable. This leaves an open question for future research, i.e., how cognitive and neural mechanisms underlying decision-making resulting in civilian casualties during intergroup conflict vary along escalated conflict expenditures.

Second, civilian casualties in real-world situations usually take place during intergroup conflict with a history. People make decisions to attack the outgroup by considering not only the consequence of the upcoming attack on the outgroup but also potential feedback/revenge from the outgroup that may occur to the ingroup thereafter. Recent theories of motivations indeed emphasize key functional roles of updated beliefs, beliefs of others, and plans of action in decision making^53^. Our paradigm modeled an initial decision making in response to intergroup conflict without providing the participants with the information of potential feedback/revenge from the outgroup. While there are indeed similar situations in real-world intergroup conflict, future research on civilian casualties during intergroup conflict should consider decision making in complicated situations when decision makers receive feedback or potential revenge from the outgroup after each decision making.

Third, our work recruited three cultural samples (i.e., Chinese, Jewish and Palestinian participants) who showed similar decreased harm avoidance related to outgroup noncombatants as conflicts escalated but difference in harm preference for outgroup combatants. It is important to test other cultural samples to verify whether collateral damage arising from intergroup conflict has similar cognitive and neural underpinnings in other sociocultural contexts. This is necessary given broad sociocultural differences in^54–56^ and potential interactions of biological and cultural factors^57–59^ on cognition and brain underpinnings of human behavior.

Finally, even though we showed evidence for an association between decreased harm avoidance and inhibition of the left MFG activity during escalation of intergroup conflict, it remains unclear whether the left MFG activity is necessary for controlling civilian casualties during intergroup conflict. This can be clarified in future research by integrating the paradigm developed in our work with the technique for interrupting brain activity such as transcranial magnetic stimulation^60^.

In conclusion, by integrating brain imaging and statistical modeling of punishment decision-making during intergroup conflict in three cultural samples, the current work disentangled harm preference and harm avoidance as two psychological constructs underlying punishment decision making related to outgroup combatants versus outgroup noncombatants. In addition, our fMRI results suggest different neural circuits supporting these two psychological constructs. Most important, we showed evidence that avoidance of harming outgroup noncombatants decreased when intergroup conflict escalated and that the decreased harm avoidance was associated with inhibition of the left middle frontal activity during selection of punishment decisions. These findings shed light on the neurocognitive underpinnings of decision-making involving civilian casualties during intergroup conflict.

## Methods

### Participants

Sample sizes were pre-determined using G*Power^61^. In Experiment 1, we aimed to conducted one-sample t-tests for each parameter in the logistic regression model and two-sample t-tests to compared parameters between the group and individual conflict conditions. Based on G*Power estimation, a sample size of 54 participants was required to obtain a medium effect size of 0.5 with an error probability of 0.05 and power of 0.95 in two-tailed one-sample t-tests. For the two-tailed two-sample t-tests, a sample size of 105 participants for both the group and individual conflict conditions were required to obtain a medium effect size of 0.5 with an error probability of 0.05 and power of 0.95. The sample size of the Jewish and Palestinian groups in Experiment 2 was determined to be similar with those in the group and individual conflict in Experiment 1. The sample size in Experiment 3 was smaller than that in Experiment 1 by considering the cost of fMRI scanning. Based on G*Power estimation, a sample size of 34 participants was required to obtain a medium effect size of 0.5 with an error probability of 0.05 and power of 0.80 in two-tailed one-sample t-tests. For the two-tailed two-sample t-tests, a sample size of 64 participants for both the group and individual conflict conditions were required to obtain a medium effect size of 0.5 with an error probability of 0.05 and power of 0.80.

In Experiment 1 we recruited 120 Chinese university students (half male, mean age ± s.d. = 22.85 ± 2.46 years) in the group conflict condition and 120 Chinese university students (half male, 21.68 ± 2.43 years) in the individual conflict condition. In Experiment 2 we recruited 106 Israeli-Jewish university students (46 males, 24.97 ± 3.78 years). To assure that only Arab participants who clearly identify themselves as Palestinian participated in our study, we originally recruited 152 Arab-Palestinian participants who completed a scale in which they rated their ethnic identification from 0% Palestinian to 100% Palestinian. Only participants who rated themselves as 100% Palestinian were further invited to participate in Experiment 2 (N=106, 46 males, 21.82 ± 2.23 years). During recruitment we did not reveal to participants that our study examined questions related to intergroup conflict so as to minimize any effects of participants’ prior attitudes about the Israeli-Palestinian conflict on behavioral measurements. All participants in Experiment 2 (i.e., both Jewish and Palestinian participants) were Israeli citizens.

In Experiment 3 we recruited 64 Chinese university students (half males, 23.13 ± 1.82 years) in the group conflict condition. Four participants were excluded due to excessive head motion during scanning, leaving 29 combatants (14 males, 22.93 ± 1.56 years) and 31 noncombatants (16 males, 23.19 ± 2.06 years) for analyses of data in the group conflict condition. An independent group of 64 Chinese university students (half males, 22.48 ± 2.42 years) were recruited in the individual conflict condition. Five participants were excluded due to excessive head motion during scanning, leaving 29 combatants (14 males, 22.76 ± 2.84 years) and 30 noncombatants (15 males, 22.23 ± 2.08 years) for analyses of data in the individual conflict condition. All participants had normal or corrected-to-normal vision and reported no neurological or psychiatric history. Informed consent was obtained prior to the experiment. All participants were paid for their participation. Experiments 1 and 3 were approved by the local ethics committee at the School of Psychological and Cognitive Sciences (Reference number: 2105-12-04), Peking University. Experiment 2 was approved by the local ethics committee at the Rambam Medical Center, Haifa University (Reference number: 044/17).

### Experiment Procedure

On each testing day, 4 participants of the same gender were recruited for a testing group who had not known each other before their participation. The experimental procedures consisted of four phases.

#### Phase 1: Group formation

In the group conflict condition, participants were randomly assigned to two groups of 2 persons by choosing a card. Participants from each group were asked to wear T-shirts of the same color (red or blue). Participant introduced their own names, nicknames, majors, and hobbies to get familiar with each other. They then started to play the Saboteur card game (http://www.annarbor.com/entertainment/saboteur-card-game-review/). During this game ingroup members played cards to build a tunnel to a gold location or to block the tunnel to prevent outgroup members from reaching the location. This game required ingroup members to cooperate with each other but to interrupt outgroup members so as to reach the gold location before the outgroup. The intergroup relationship was built by playing this game for 50 minutes. To check the effectiveness of the group manipulation, after the game, participants were asked to complete a modified version of the Inclusion of Other in the Self Scale^19^ to assess their feelings of closeness between oneself and ingroup members and between oneself and outgroup members. The individual conflict condition was the same as the group conflict condition except that participants were not divided into groups and they played the Saboteur card game against the other three participants. After the game, participants were asked to assess their feelings of closeness between oneself and the other three participants with a modified version of the Inclusion of Other in the Self Scale^19^. Phase 1 lasted for 60 minutes.

#### Phase 2: Inducing conflict

For the group conflict condition one participant from each group was randomly assigned as a combatant who allotted 100 RMB (in Experiment 1; 100 Shekels in Experiment 2; and 200 RMB in Experiment 3) between the two combatants by giving different weights to a number of plans. The other participant from each group waited, as a noncombatant, while combatants made their decisions (Phase 2 in Figure 1). This procedure was the same for the individual conflict condition except that 2 participants were randomly assigned as combatants and 2 as noncombatants.

There were nine different money distribution plans, which included Self(55)/Rival(45), Self(60)/Rival(40), Self(65)/Rival(35), Self(70)/Rival(30), Self(75)/Rival(25), Self(80)/Rival(20), Self(85)/Rival(15), Self(90)/Rival(10), Self(95)/Rival(5) (The ratios of money allocation plans were the same across three experiments). Each combatant was given 108 cards and had to allot these cards to the nine plans (there must be at least 1 card for each plan). Both combatants and noncombatants were informed that a computer would randomly select one out of 108 cards, which corresponded to one money allocation plan from each combatant. That is, a plan with more cards allocated would be more likely to be selected by the computer. After the computer selected a plan from each combatant, the money would be finally divided between the two combatants according to the plan with a larger difference between Self and Rival. If two plans from the two combatants were the same, the computer would randomly select one plan to decide the final payment. There were typically two strategies to allocate the cards. Allotting more cards to Self(55)/Rival(45) plan represents a fairness strategy whereas more cards to Self(95)/Rival(5) represents a selfish strategy. All plans allot more money to self and the rules benefited the combatant who employed a more selfish strategy. These designs induced conflict between two combatants. The two combatants finished the money allocation task in two private rooms so that the other participants would not know their decisions. All the participants were informed that each noncombatant would receive 100 RMB for their participation. Each combatant, however, would received a basic payment of 50 RMB and share a 100 RMB bonus with the other combatant based on the final money allocation plan. On average each combatant would received 100 RMB for his/her participation. Phase 2 lasted for 15 minutes.

#### Phase 3: Pain threshold measurement

To help participants to have an intuitive feeling of different levels of electric shocks, each participant underwent a pain titration procedure with a Digitimer DS7 electric stimulator individually (Phase 3 in Figure 1). Following a brief overview of the equipment and titration process, two electrodes were attached to the back of the left hand. The titration procedure began with a low-voltage electric shock (0.5 mA). A participant was then asked to rate his/her subjective feelings of pain on an 11-point scale (0 = no pain, 10 = intolerable). The initial rating was followed by a series of shocks increasing with a step of 0.5 mA. Subjective ratings of pain were collected after each shock until a rating of 10 was reached. The same procedure was repeated twice for each participant. Phase 3 lasted for 40 minutes.

#### Phase 4: Punishment decision making

In this phase, the participants were informed that the 108 money allocation plans from one combatant would be presented one by one. There were 108 trials with equal numbers of the nine money distribution plans which were actually created by the computer program and randomly presented to participants. On each trial, a money distribution plan was first presented for 4 s on a screen. Thereafter, two options of electric shocks were shown on the left and right side of the screen, respectively (Phase 4 in Figure 1). Each option (either the left or right) consisted of two vertical bars that indicated intensities of two electric shocks. The shock intensities were indexed by rating scores of subjective painful feelings (from 1 = slightly pain to 9 = extremely painful) that each participant experienced in Phase 3. In the group conflict condition, one shock would be applied to the outgroup combatant as punishment and the other to the outgroup noncombatant as a “collateral damage”. In the individual conflict condition, one shock would be applied to the combatant as punishment and the other to a bystander. The intensities of two electric shocks in one option for punishing the combatant and harming the noncombatant, the intensities of two electric shocks in the left and right options, and the intensities of electric shocks varied independently across trials. Non-independent variations between the shock intensities might lead to high collinearity between two variables and result in unstable estimation of the regression parameters. The participants were asked to make a choice between the left and right options by pressing one of two buttons using the right index and middle fingers. A fixation of 2 s was presented after a button press when the next trial started. They were also informed that electric shocks in one of their decisions would be randomly chosen to be applied to the target combatant and noncombatant after the experiment, which, though were not really applied to the participants who were debriefed. All the four participants in each session knew that the two outgroup members would make decisions that might lead to them receiving electric shocks. C1/C2 and N1/N2 performed the decision making task simultaneously in private rooms during Phase 4.

Task instructions in Phases 2-4 were given at the beginning of Phase 2 to all the participants simultaneously. The combatants and the noncombatants knew each other’s tasks. That means, N1/N2 knew that they would be subject to electric shocks when the other participants made decisions to punish the combatants. All the participants in Phase 4 knew that other participants were doing the same decision-making tasks.

According to the unfairness of the money allocation plan, all 108 trials were coded as being in one of three conflict levels. In low, middle and high conflict levels, the target combatant would keep 55 – 65 RMB, 70 – 80 RMB and 85 – 95 RMB for oneself, respectively. In each conflict level, there were 36 trials with 36 combinations of two options of electric shocks. Specifically, nine levels of electric shocks were divided into three pain levels (e.g., low, middle and high pain). These three pain levels towards combatant and noncombatant would create 9 different combinations of options. In each trial, two out of nine options were selected and presented. This created 36 different combinations of two options. To this end, the difference of electric shocks between two options will vary in each trial and vary independently between combatant and noncombatant, which were entered into the following logistic regression modelling. Finally, the experiment design and purpose were explained in debriefing and all participants received 100 RMB for participation. Phase 4 lasted for 35 minutes.

The experimental procedures of Experiment 2 in Jewish and Palestinian participants were the same as those in Study 1 except the following. There was no group formation phase at the start. Instead, two Jewish participants and two Palestinians (all were of the same gender and were strangers to each other) were recruited each time and they introduced their nationality to each other. Furthermore, in order to highlight the nationality of each participant, participants of the same nationality performed a priming task together which contained images of items that relate to one’s nationality. Such as the symbols of each group’s nation (e.g. flags, emblems, anthems), which was found to increase the feeling of identification with one’s own nation^27^, as well as photos of famous people (e.g., Yasser Arafat vs. Shimon Peres) and holy places (e.g., Al-Aqsa Mosque vs. Hakotel). The participants from each group were asked to jointly rate the feelings of belonging and excitement that each image evoked (1 = very little, 9 = very much).

The experimental procedures of Study 3 in the neuroimaging lab were the same as those in Study 1 except the following. The Saboteur card game lasted 90 minutes during Phase 1 to strengthen the intergroup relationship since the whole experiment lasted for a longer time in the Experiment 3. Each participant was scanned using fMRI while making punishment choices during Phase 4. There were 108 trials for each participant which were equally distributed into six scans. In each scan, there were 18 trials which included 6 low conflict trials, 6 middle conflict trials, and 6 high conflict trials. On each trial, a jitter fixation was first presented for 2, 4, 6, 8, or 10 s. A money distribution option given by the target combatant was then presented for 2 s. A following jitter fixation was presented for 2, 4, 6, 8, or 10 s. Thereafter, two options of electric shocks towards the target combatant and noncombatant were presented and participants were asked to make a choice between the options within 5 seconds. Each scan started with a 12-s fixation to get a baseline for blood-oxygen-level-dependent (BOLD) signals. The last trial of each scan was followed by a 12-s fixation.

### Model comparisons

We tested additional two models (Models 2 and 3) and compared these models with the one we reported in the main text (Model 1). Model 2 only included intercept in the logistic regression model to regress the inherent tendency to choose left or right option of punishment decisions for each participant (ΔV = intercept; the participants tended to choose one (the left or right) option regardless of shock intensities and monetary distribution plans). This is a baseline control model compared with other models. Model 3 hypothesized that outgroup combatants and noncombatants are treated in the same way during intergroup conflict. We included two independent variables in the regression model to account for the probability that participants would select the left (*p*) or right options (*1-p*) in each trial. These included the difference in the sum of shock intensities to the combatant and the noncombatant between left and right options (ΔS = ΔS_c_ + ΔS_n_) and the interaction between ΔS and level of conflict (L). The logistic regression model was defined as following:

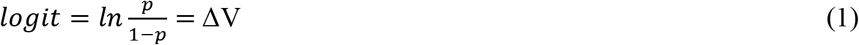

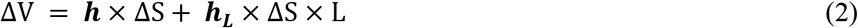

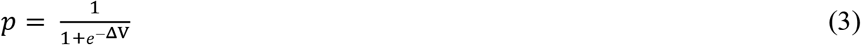

A larger *p* is associated with higher possibility of selection of the left option.A positive ***h*** value indicates harm preference towards both combatants and noncombatants — a propensity to select an electric shock option with a greater sum of intensities from two options. A negative ***h*** value indicates harm avoidance towards both combatants and noncombatants — a propensity to select an electric shock option with a smaller sum of intensities from two options. ***h_L_*** manifest how ***h*** varies along escalated levels of conflict (positive and negative ***h_L_*** value indicates increase or decrease of ***h***, respectively).

We compared these models using Bayesian Information Criterion (BIC), prediction accuracy, and area under the curve (AUC). BIC penalizes models with a greater number of parameters, and the model with the lowest group BIC is the preferred model. We computed BIC scores for each model fit in each individual participant and summed the BIC scores across subjects to obtain a group BIC score. We also compared predicted choice probability of each model with the categorization threshold of 0.5 and participants’ choices in all trials to calculate the prediction accuracy of each model. If the predicted choice probability is larger than 0.5 in one trial, the model will predict that participant choose left option in that trial. Otherwise, participant will choose right option. We calculated mean prediction accuracy across all participants for each model. The model with the highest prediction accuracy is the preferred model. AUC was calculated as the area under the Receiver Operator Characteristic curve, which provides a simple way to summarize all the confusion matrices by setting different categorization thresholds. The model with the largest AUC is the preferred model.

### fMRI data acquisition and preprocessing

Brain images were acquired using a 3.0T GE Signa MR750 scanner (GE Healthcare; Waukesha, WI) with a standard 8 channel head coil. Functional images were acquired by using T2-weighted, gradient-echo, echo-planar imaging (EPI) sequences sensitive to BOLD signals (64 × 64 × 32 matrix with 3.75 × 3.75 × 5 mm^3^ spatial resolution, repetition time = 2000 ms, echo time = 30 ms, flip angle = 90°, field of view = 24 × 24 cm^2^). A high-resolution T1-weighted structural image (512 × 512 × 180 matrix with a spatial resolution of 0.47 × 0.47 × 1.0 mm^3^, repetition time = 8.204ms, echo time = 3.22ms, flip angle = 12°) was acquired after the first three functional scans. Padded clamps were used to minimize head motion and earplugs were used to attenuate scanner noise. The stimuli were projected onto a screen at the head of the magnet bore using Presentation. Participants viewed the screen through a mirror attached to the head coil.

Functional images were preprocessed using SPM12 software (the Wellcome Trust Centre for Neuroimaging, London, UK). Head movements were corrected within each run and six movement parameters (translation; x, y, z and rotation; pitch, roll, yaw) were extracted for further analysis in the statistical model. The anatomical image was coregistered with the mean realigned functional image. The functional images were resampled to 3 × 3 × 3 mm^3^ voxels, normalized to Montreal Neurological Institute (MNI) template space and then spatially smoothed using an isotropic of 8 mm full-width half-maximum (FWHM) Gaussian kernel.

### Whole brain parametric analyses

The first level general linear model (GLM) was constructed to identify brain regions responding parametrically at choice onset to the difference of electric shocks between chosen and unchosen options in each participant. The GLM contained three sets of event regressors describing: (1) the onsets of the money allocation plans, the corresponding duration of 1 TR, and a parametric modulator of different money allocation plans representing three different levels of conflict (L); (2) the onsets of two options of shocks, the corresponding duration of participants' reaction time on each trial, and four parametric modulators: (i) the difference of electric shocks between chosen and unchosen options towards combatant (ΔS_c_); (ii) the difference of electric shocks between chosen and unchosen options towards noncombatant (ΔS_n_); (iii) the interaction between conflict level and difference of electric shocks towards combatant (ΔS_c_ * L); (iv) the interaction between conflict level and difference of electric shocks towards noncombatant (ΔS_n_ * L); (3) the onsets of the button press for punishment decision making and a duration of zero. Besides, the GLM also included the realignment parameters to account for any residual movement-related effect.

### Whole brain regression analyses

We conducted four second-level whole-brain one-sample t-tests with parameters estimated by the logistic regression modeling as covariates. These tests were constructed to examine brain regions associated with individual differences in harm tendency (***h_c_***), harm avoidance (***h_n_***), variation of harm tendency with conflict levels (***h_c-L_***), and variation of harm avoidance with conflict levels (***h_n-L_***). The first regression analysis included the contrast images of parametric modulation effects of ΔS_c_ with ***h_c_*** as the covariate. The second regression analysis included the contrast images of parametric modulation effects of ΔS_n_ with ***h_n_*** as the covariate. The third regression analysis included the contrast images of parametric modulation effects of ΔS_c_ * L with ***h_c-L_*** as the covariate. The fourth regression analysis included the contrast images of parametric modulation effects of ΔS_n_ * L with ***h_n-L_*** as the covariate. The brain regions associated with individual differences in parameters were estimated by calculating one-sample t-test using the contrasts of [0 1] or [0 −1] to assess the positive or negative associations. Brain activations were defined by combining a whole-brain voxel-level threshold of *P* < 0.001 and a cluster-level threshold at *P* < 0.05 FWE corrected.

### Moderation analyses

To test whether the associations between ***h_n-L_*** and the left MFG activity were significantly different between the group conflict and individual conflict conditions, we conducted a second-level whole-brain two-sample t-test with ***h_n-L_*** in these two conditions as covariates. The design matrix for this t-test included the contrast images of the parametric modulation effect of ΔS_n_ * L in each condition and ***h_n-L_*** in the two conditions as the first and second covariates. The moderation effect was estimated by calculating two-sample t-test using the contrasts of [0 0 −1 1] to assess the enhanced coupling between ***h_n-L_*** and brain activity that was stronger in the group conflict condition than in the individual conflict condition. Small volume corrections within the mask of the left MFG defined using the Brainnetome Atlas (Region 22: MFG_L_7_4_9 (46v)^28^ were applied to search for significant results with the threshold of p<0.05, FWE corrected for each voxel.

### Bayesian factor analyses

Bayesian factor analyses were used to compare the likelihood of the data fitting under the null hypothesis with the likelihood of fitting under the alternative hypothesis^23^. Compared with widely used p-values, the Bayes factor allows the researcher to make a statement about the alternative hypothesis, rather than just the null hypothesis. It also provides a clearer estimate of the amount of evidence present in the data. In the current paper, Bayes factor was calculated using the r toolbox “Bayesian factor” (https://richarddmorey.github.io/BayesFactor/). BF_01_ represents the ratio that contrasts the likelihood of the data fitting under the null hypothesis with the likelihood of fitting under the alternative hypothesis. On the contrary, BF_10_ represents the ratio that contrasts the likelihood of the data fitting under the alternative hypothesis with the likelihood of fitting under the null hypothesis. Therefore, as BF_01_ increases, there is more evidence in support of the null hypothesis. While as BF_10_ increases, there is more evidence in support of the alternative hypothesis. A Bayesian factor falling between 1 to 3 is regarded as weak evidence, between 3 to 20 is regarded as positive evidence, between 20 to 150 is regarded as strong evidence and larger than 150 is regarded as very strong evidence in support of null (BF_01_) or alternative hypothesis (BF_10_)^62^.

## Data availability

The data that support the findings of this study are available from the corresponding author upon reasonable request.

## Code availability

The codes used to analyze the data that support the findings of this study are available from the corresponding author upon reasonable request.

## Acknowledgements

This research was supported by the National Natural Science Foundation of China (projects 31421003, 31871134, and 31661143039), the Ministry of Science and Technology of China (2019YFA0707103), and the Israel Science Foundation (2510/16). We thank the National Center for Protein Sciences at Peking University for assistance with this study. The authors thank L. Zhu, J. Li, and Y. Ma for insightful discussion about experimental design and data analyses, R. Hampton for proof reading, and S. Wang for modification of Figure 1. The funder had no role in the conceptualization, design, data collection, analysis, decision to publish or preparation of the manuscript.

## Author contributions

X. H. and S.H. conceived the research programme and designed the experiments. X. H., S. Z., N. F., T. W., T. G., S. S., X. W., S. H. carried out the experiments. X. H., N. F., and S. H. analyzed the data. X. H., S. H., and S. S. wrote the paper. M.J.G reviewed and edited the paper. S.H. supervised the entire work.

## Additional information

Supplementary information is available for this paper.

