## Supplementary information for "Cognitive and neural bases of decision-making causing civilian casualties during intergroup conflict"

### Supplementary Results

#### Result 1: Statistical details of behavioral results in Experiment 2

Because of the difference in national and religious identity between the two ethnic groups in Experiment 2, we divided 2 Jewish participants and 2 Palestinian participants (all were of the same gender and strangers) into two mini-groups on each testing session. To highlight participants' national identity, after introducing one's own nationality, all participants were presented with images of items or celebrities/holy places that remind them of the nationality of Israel or Palestine. The participants from each mini-group were asked to jointly observe each picture and rate their feelings of picture-induced belonging and excitement (1 = very little, 9 = very much). The rating scores were high for both ethnic groups and higher in Palestinian participants than in Jewish participants (see Supplementary Table 2).

#### Result 2: Results of the mediation analysis in Experiment 2.

Because $\boldsymbol{h_c}$ was significantly smaller in Jewish compared with Palestinian participants, we further explored possible mediation factors. As compromise attitudes towards intergroup conflict are associated with positive emotions towards outgroup, such as liking and understanding[63] and decreasing of negative emotion such as anger[64], we asked the participants in Experiment 2 to rate their emotions, including liking, understanding, and anger (from 1="not at all" to 5="very much") toward the other ethnic group. Relative to Palestinian participants, Jewish participants reported more liking (mean ± s.d. = 2.83±1.03 vs. 2.46±0.99, t(209) = 2.63, $P$ = 0.014, FDR corrected, Cohen's $d$ = 0.36, 95% CI = 0.09, 0.64; $BF_{10}$ = 3.76), more understanding (3.25±1.11 vs. 2.92±1.02, t(209) = 2.20, $P$ = 0.029, FDR corrected, Cohen's $d$ = 0.30, 95% CI = 0.03, 0.61; $BF_{10}$ = 1.44), and less anger (2.42±1.20 vs. 3.02±1.23, t(209) = -3.58, $P$ = 0.001, FDR corrected, Cohen's $d$ = 0.49, 95% CI = -0.93, -0.27; $BF_{10}$ = 54.34). We further conducted mediation analyses to examine possible mediation roles of these emotions. The results showed, among the three emotions, liking partially mediated the difference in $\boldsymbol{h_c}$ between Jewish and Palestinian participants (The indirect effect size was -0.05 with a 95% confidence interval which did not include zero (-0.10, -0.01); see Supplementary Fig. 3 and Supplementary Table 3 for statistical details).

**Result 3: Statistical details of behavioral results in Experiment 3**

One sample t-tests showed that $\boldsymbol{h_c}$ did not differ significantly from zero in either group conflict condition or individual conflict condition (see Supplementary Table 4 and Supplementary Fig. 6 for statistical details). There was no significant difference in $\boldsymbol{h_c}$ between the group and individual conflict conditions. The results provide no evidence that harm preference towards outgroup combatants influence punishment decisions on average across all conflict levels. However, $\boldsymbol{h_n}$ was significantly smaller than zero in both the group conflict condition and individual conflict condition. Moreover, $\boldsymbol{h_n}$ was significantly less negative in the group conflict condition than in the individual conflict condition, suggesting that the distribution of a larger amount of money in Experiment 3 (200 RMB) compared to those in Experiment 1 (100 RMB) might result in higher levels of conflict and lead to less harm avoidance in the group conflict condition.

Similarly, the analyses of $\boldsymbol{h_{c\text{-}L}}$ or $\boldsymbol{h_{n\text{-}L}}$ revealed different patterns in the group and individual conflict conditions. $\boldsymbol{h_{c\text{-}L}}$ was significantly larger than zero in both the group and individual conflict conditions, but did not differ significantly between the group and individual conflict conditions. The results suggest that, for both the group and individual conflict conditions, harm preference towards combatant increased as the level of conflict escalated. Nevertheless, $\boldsymbol{h_{n\text{-}L}}$ was significantly larger than zero in the group conflict condition but not in the individual conflict condition. There was a significant difference in $\boldsymbol{h_{n\text{-}L}}$ between the group and individual conflict conditions. These results replicate the findings in Experiments 1 and 2 and provide further evidence that decreased harm avoidance served as a cognitive basis of collateral damage as intergroup conflict escalated.

In Experiment 3 the logistic regression model also correctly predicted a high proportion of punishment decisions across trials, which are far beyond the chance level (81.88%±7.50%, $t(135) = 49.54$, $P < 0.001$, Cohen's $d = 4.25$, 95% CI = 80.60%, 83.15%, $BF_{10} > 150$, see Supplementary Fig. 1).

### Supplementary Tables

**Supplementary Table 1:** Results of model comparisons in all experiments.

| Experiment | Model | BIC | Prediction accuracy (Mean±SD) | AUC (Mean±SD) |
|---|---|---|---|---|
| 1 | Model 1 | **18159.58** | **0.85±0.08** | **0.91±0.08** |
| | Model 2 | 35584.88 | 0.54±0.11 | 0.50±0.00 |
| | Model 3 | 27061.69 | 0.70±0.11 | 0.77±0.12 |
| 2 | Model 1 | **17076.65** | **0.84±0.08** | **0.89±0.09** |
| | Model 2 | 31442.07 | 0.56±0.05 | 0.50±0.00 |
| | Model 3 | 21669.25 | 0.77±0.10 | 0.81±0.12 |
| 3 | Model 1 | **10456.98** | **0.82±0.08** | **0.88±0.07** |
| | Model 2 | 20170.1 | 0.52±0.17 | 0.50±0.00 |
| | Model 3 | 15231.57 | 0.72±0.09 | 0.78±0.11 |

BIC=Bayesian Information Criterion; AUC=area under the curve. Smaller BIC but larger prediction accuracy and AUC indicate better model fit. All the results of BIC, prediction accuracy, and AUC support that Model 1 is the best model of the participants' punishment decisions across three Experiments.

**Supplementary Table 2:** Rating scores (mean ± s.d.) of manipulation check in Experiment 2.

| | Jewish participants | Palestinian participants | Jewish vs. Palestinian participants | | | | |
|---|---|---|---|---|---|---|---|
| | | | **t** | ***P*** | **d** | **95%CI** | **$BF_{10}$** |
| Belonging | 6.44±1.68 | 7.04±1.37 | 2.05 | 0.042 | 0.38 | 0.02, 1.17 | 1.31 |
| Excitement | 5.96±1.62 | 6.88±1.16 | 3.45 | 0.001 | 0.65 | 0.39, 1.45 | 35.54 |

Note: Rating score: 1 = very little, 9 = very much; two tailed t-tests compared rating scores of the ethnic groups.

**Supplementary Table 3:** The results of the mediation analysis in Experiment 2.

| Variable | Coeff | SE | t | *P* |
|---|---|---|---|---|
| **Regression Model 1 (Total effect of Ethics on $h_c$)** | | | | |
| Independent: Ethics | -0.250 | 0.073 | -3.412 | <0.001 |
| Dependent: $h_c$ | | | | |
| **Regression Model 2 (Ethics to Liking)** | | | | |
| Independent: Ethics | 0.366 | 0.139 | 2.634 | 0.009 |
| Mediator: Liking | | | | |
| **Direct effect of Liking on $h_c$** | | | | |
| Mediator: Liking | -0.135 | 0.035 | -3.821 | <0.001 |
| Dependent: $h_c$ | | | | |
| **Remaining direct effect of Ethics on $h_c$** | | | | |
| Independent: Ethics | -0.201 | 0.072 | -2.779 | 0.006 |
| Dependent: $h_c$ | | | | |
| | ***Coeff*** | ***SE*** | ***LLCI95*** | ***ULCI95*** |
| **Indirect effect of Ethics on $h_c$ via Liking (bootstrap result)** | | | | |
| Liking | -0.049 | 0.022 | -0.098 | -0.011 |

Notes. Confidence intervals for indirect effect are bias-corrected and accelerated; bootstrap resamples = 5000; N = 211 for all tests.

**Supplementary Table 4:** Group-level statistical details of logistic regression parameters in Experiment 3.

| | **Mean (s.d.)** | **t** | **p** | **Cohen's d** | **95% CI** | **$BF_{10}$** |
|---|---|---|---|---|---|---|
| Group conflict condition | | | | | | |
| $h_c$ | 0.18 (0.77) | 1.79 | 0.078 | 0.23 | [-0.02, 0.38] | 0.63 |
| $h_n$ | -0.28 (0.27) | -8.04 | <0.001 | 1.04 | [-0.35, -0.21] | >150 |
| $h_{c\text{-}L}$ | 0.36 (0.54) | 5.20 | <0.001 | 0.67 | [0.22, 0.50] | >150 |
| $h_{n\text{-}L}$ | 0.12(0.18) | 5.36 | <0.001 | 0.69 | [0.08, 0.17] | >150 |
| Individual conflict condition | | | | | | |
| $h_c$ | -0.01 (0.43) | -0.10 | 0.919 | 0.01 | [-0.12, 0.11] | 0.14 |
| $h_n$ | -0.50 (0.49) | -7.83 | <0.001 | 1.02 | [-0.62, -0.37] | >150 |
| $h_{c\text{-}L}$ | 0.27 (0.28) | 7.25 | <0.001 | 0.94 | [0.19, 0.34] | >150 |
| $h_{n\text{-}L}$ | 0.02 (0.19) | 0.94 | 0.467 | 0.12 | [-0.03, 0.07] | 0.22 |
| Differences between group and individual conflict conditions | | | | | | |
| $h_c$ | | 1.61 | 0.148 | 0.29 | [-0.04, 0.41] | 0.62 |
| $h_n$ | | 3.06 | 0.008 | 0.56 | [-0.36, -0.08] | 12.19 |
| $h_{c\text{-}L}$ | | 1.19 | 0.236 | 0.22 | [-0.06, 0.25] | 0.37 |
| $h_{n\text{-}L}$ | | 2.97 | 0.008 | 0.54 | [0.03, 0.17] | 9.53 |

### Supplementary Figures

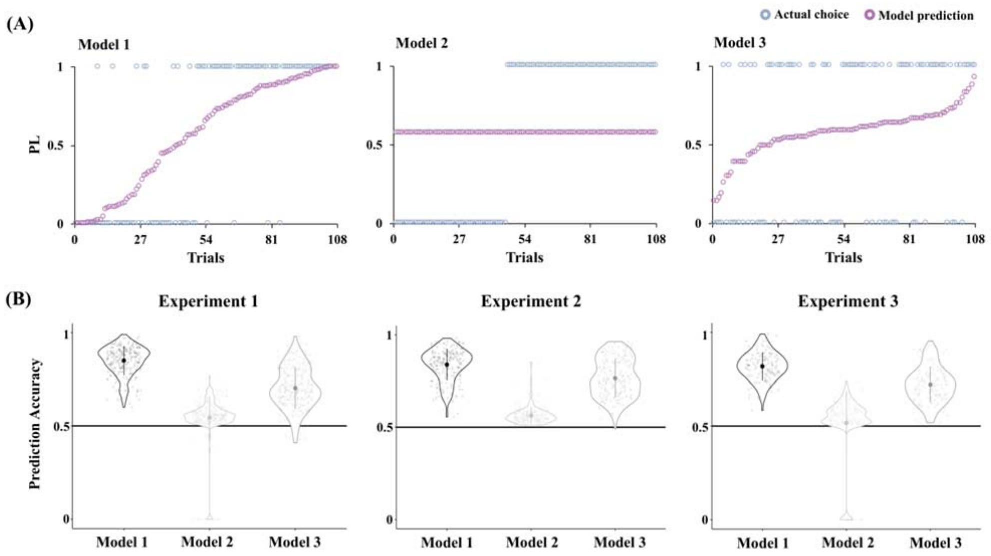

**Supplementary Fig. 1:** Prediction accuracies of different models of participants' choices of electric shocks. (A) An example data from a participant in the group conflict condition in Experiment 1. *p* represents probability to choose the left option. If $p > 0.5$, the model predicts selection of the left option; otherwise, the model predicts selection of the right option. The violet circle represents the probability of choosing the left option by the corresponding models. The blue circle represents a participant's real choice in that trial. A blue circle with a value of zero along y-axis indicates a choice of the right option whereas a value of 1 indicates a choice of the left option. (B) Prediction accuracies of the three testing models. The prediction accuracy of Model 1 is the best, and Model 2 is the worst. Violin plots show the distributions of prediction accuracy with the large dots showing the means of the prediction accuracy and a vertical bar indicating standard deviation of the prediction accuracy. Each small gray dot represents one participant. see Methods for details of the three testing models.

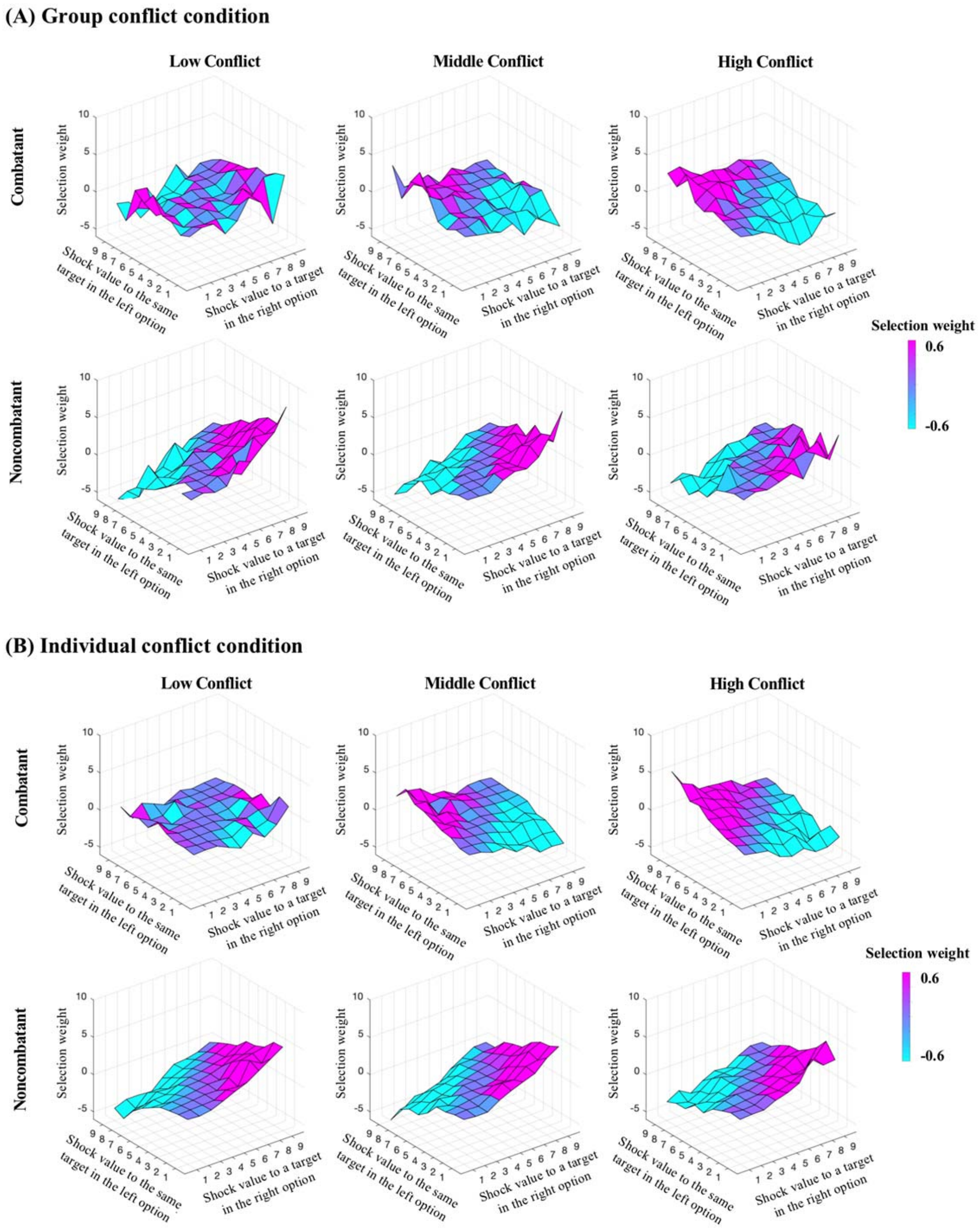

**Supplementary Fig. 2:** Illustration of selection weight of the left and right options corresponding to specific shock intensities shown in Phase 4 in Experiment 1. Selection weight was calculated as: $\text{Selection weight} = \frac{1}{n} * \sum_{i=1}^{n} \text{Choice}_i * \text{abs}(\Delta S_i)$; n = number of participants; $\Delta S_i$ = shock value to a target in the left option minus shock value to the same target in the right option of the participant *i*; $\text{Choice}_i$ = 1 if the participant *i* choosing the left option of punishment on one trial; $\text{Choice}_i$ = -1 if the participant *i* choosing the right option of punishment on one trial; abs = absolute value. Selection weight represents tendencies to select the left or right options when considering the difference in shock intensity related to combatant (or noncombatant) between the left and right options across all participants. (A) Selection weight in the group conflict condition. Figures illustrate a sharp pattern of selection of the left or right choice in correspondence with the magnitudes of shocks to outgroup combatants particularly when conflict was high. By contrast, there is a sharp pattern of selection of the left or right choice in correspondence with the magnitudes of shocks to outgroup noncombatants particularly when conflict was low. (B) Selection weight in the individual conflict condition. Similarly, figures illustrate a sharp pattern of selection of the left or right choice in correspondence with the magnitudes of shocks to outgroup combatants particularly when conflict was high. However, there are similar sharp patterns of selection of the left or right choice in correspondence with the magnitudes of shocks to outgroup noncombatants regardless conflict levels.

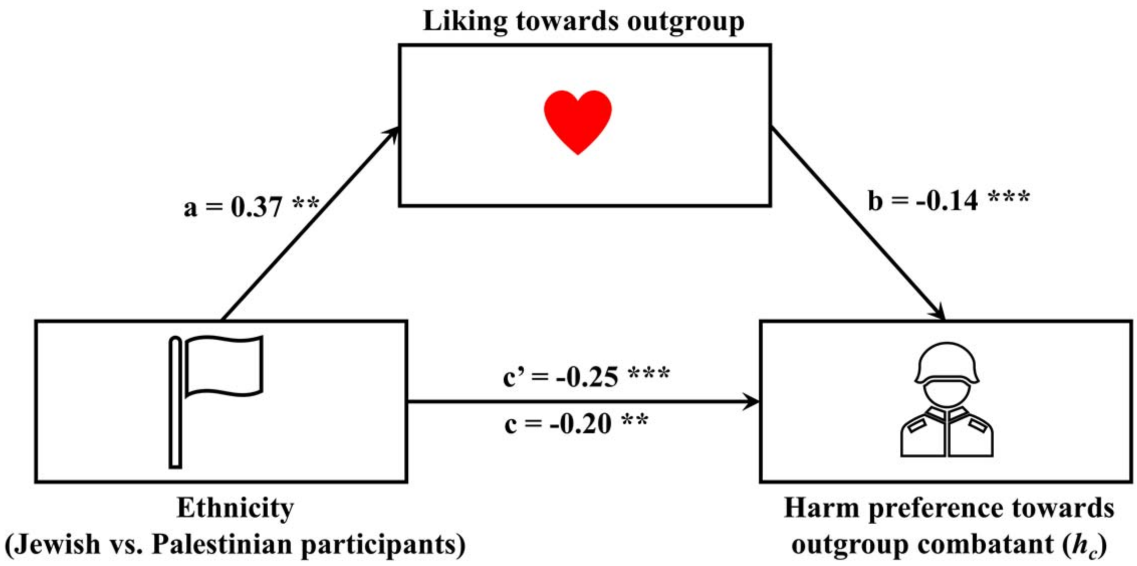

**Supplementary Fig. 3:** Results of the mediation analysis in Experiment 2. Self-report of liking for outgroup partially mediates the relationship between participants' ethnicity and harm preference towards outgroup combatant ($\boldsymbol{h_c}$).

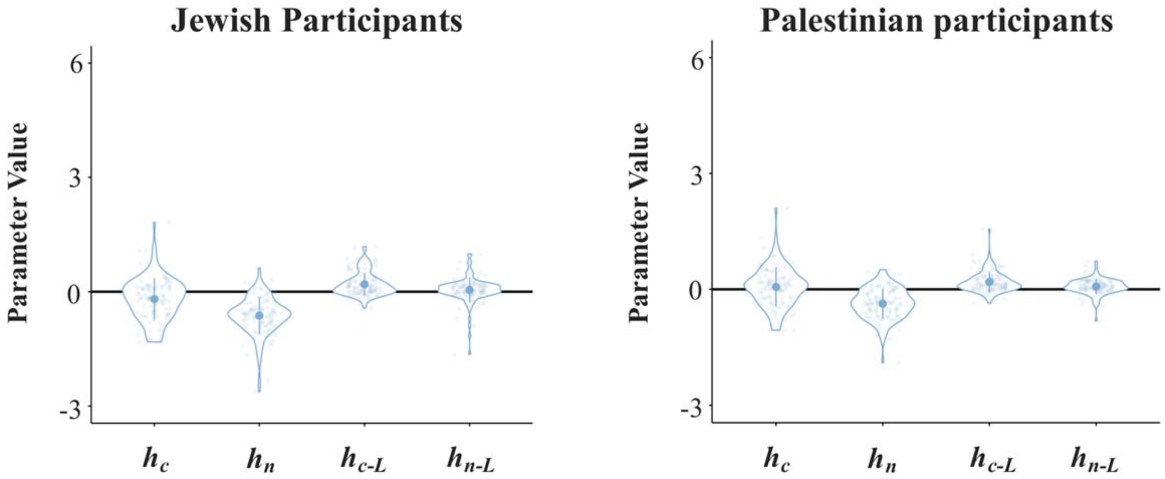

**Supplementary Fig. 4:** Separate logistic regression parameters in Jewish and Palestinian participants in Experiment 2. Violin plots show means (big dots), s.d. (bars), and distributions of parameter values. Each dot represents an individual participant.

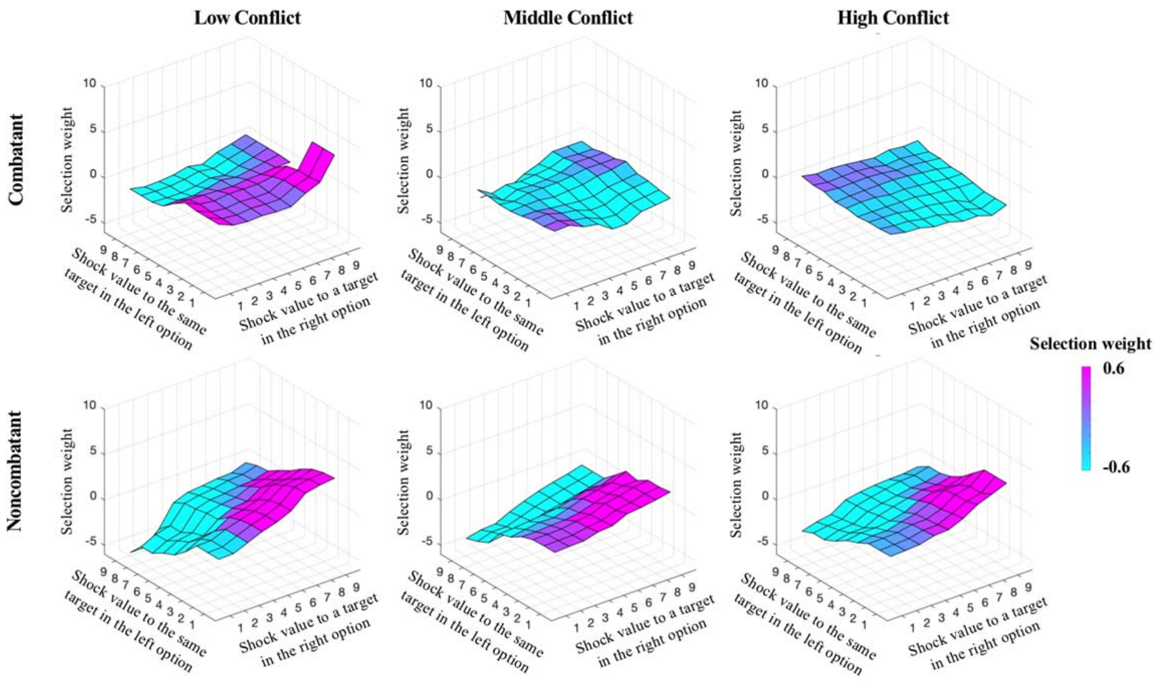

**Supplementary Fig. 5:** Illustration of selection weight of the left and right options corresponding to specific shock intensities shown in Phase 4 in Experiment 2. Selection weight is calculated in the same way as that in Experiment 1. Missing selection weights due to random generation of options of shock intensities, which did not influence the logistic regression modeling, were calculated as the average of the surrounding selection weights.

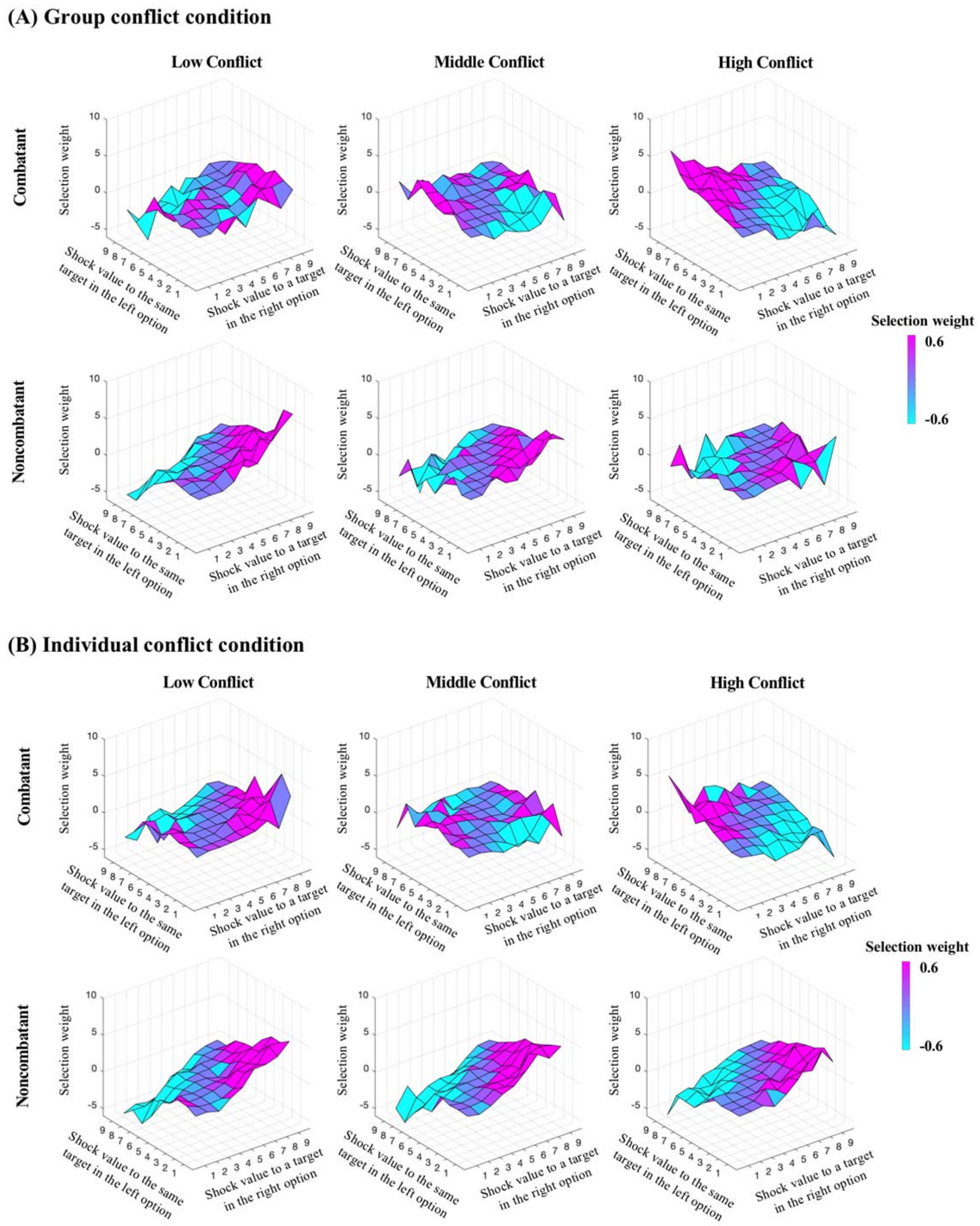

**Supplementary Fig. 6:** Illustration of selection weight of the left and right options corresponding to specific shock intensities shown in Phase 4 in Experiment 3. Selection weight is calculated in the same way as that in Experiment 1. (A) Selection weight in the group conflict condition. (B) Selection weight in the individual conflict condition.
